# DNA-Lipid Nanodiscs with a Polyethylene Glycol Interface

**DOI:** 10.64898/2026.02.16.705827

**Authors:** Soumya Chandrasekhar, Christopher Maffeo, Sanjai Karanth, Rachel Bricker, Joy Kabuga, Diana Patricia Nunes Gonçalves, Aleksei Aksimentiev, Thorsten-Lars Schmidt

**Author notes:** Corresponding Author Thorsten Schmidt - Department of Physics, Kent State University, Kent, OH, 44242, USA; Advanced Materials and Liquid Crystal Institute, Kent State University, OH, USA.

## Abstract

Nanoscale bilayer mimetics such as protein or polymer-based nanodiscs are versatile tools to study the physical chemistry of lipid bilayers or the structures and functions of membrane proteins. Here, we introduce DNA-Lipid Nanodiscs (DLNs) in which the interface between hydrophobic lipids and the charged DNA is mediated through amphiphilic poly(ethylene)glycol (PEG). For this, we modified oligonucleotides with PEG and hybridized them to a single-stranded ring to form functionalized minicircles with a well-defined diameter. The center of these minicircles can be filled with a lipid bilayer through addition of detergent-solubilized lipids followed by detergent removal. Simulations reveal that the methylene groups in PEG form dynamic interactions with the acyl chains of lipids, effectively shielding the hydrophobic mismatch. As proof of concept towards incorporation of complex membrane proteins, we inserted the biotinylated transmembrane domain of synaptobrevin into these nanodiscs and bound them to streptavidin-modified quantum dots as a marker for successful incorporation. We envision these atomically precise, modular DNA scaffolds to be widely applicable in future studies of membrane proteins and nanoscale lipid membranes.

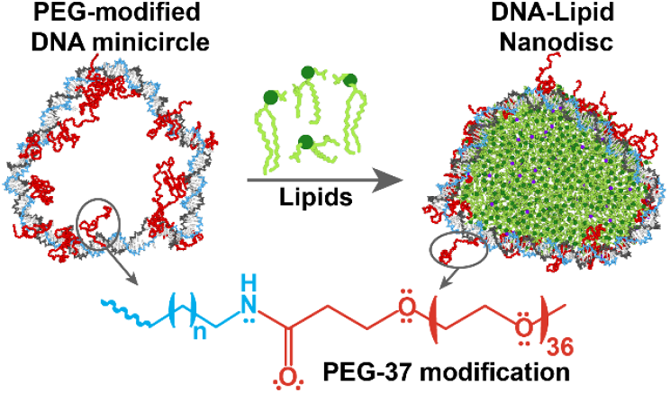

## Introduction

Lipid membranes and membrane proteins are critical components of all cells, playing vital roles in cellular structure, function and communication.^1^ Several nanoscale bilayer mimetics including liposomes,^2,3^ bicelles^4^ and nanodiscs^5–7^ have been developed to study the structure and function of membrane proteins in their native lipid environment. Nanodiscs^8^ are nanoscale, discoidal particles where the rim of the lipid bilayer is limited and stabilized by amphipathic proteins (Membrane Scaffolding Proteins - MSPs)^9^, peptides (peptidiscs),^10–12^ polymers,^13^ or S-alkyl-modified DNA.^14^ MSP nanodiscs consist of an apolipoprotein (apo) A-I belt that wraps around the lipid bilayer and typically ranges from 7-17 nm in diameter.^15^ Styrene-maleic acid (SMALPs)^13^ and various other amphiphilic copolymers such as Ultrasolute™ Amphipols^16^, poly(acrylic acid-co-styrene) (AASTY)^17^, diisobutylene-alt-maleic acid (DIBMA)^18^ can solubilize membranes.

In established bilayer mimetic systems, it is difficult to precisely predetermine the nanodisc size, to add additional functional groups or to control the position of such functionalizations on the belt. Structural DNA nanotechnology on the other hand provides an approach for the modular design and customization of nanostructures with precise control over size and the positions where functionalizations are introduced.^19^ Being a polyanion, DNA can only form interfaces with membranes by adding hydrophobic modifications to its backbone or nucleobases.^20^ Hydrophobic DNA modifications include cholesterol,^21,22^ tocopherol,^23^ lipids,^24,25^ porphyrins,^26–28^ pyrene,^29^ and S-alkylated phosphorothioates.^14,30^ Such modified DNA nanostructures were used to create DNA nanopores^30,31^ DNA-lipid nanodiscs,^14,32^ to mimic enveloped virus particles,^24^ or to deform,^33–35^ pattern,^36,37^ fuse^38,39^ and control budding^40^ of membranes. Previously, we synthesized a DNA-lipid nanodisc (DLN) from a double stranded (ds) DNA minicircle formed from a covalently closed single-stranded (ss) circle and seven identical, complementary, modified oligonucleotides, in which S-alkyl phosphorothioates enable interactions with lipid membranes.^14^ This system generated nanodiscs of uniform size which was dictated by the length of the ss circle. Subsequent coarse-grained molecular dynamics simulations suggested that greater charge neutralization could produce stronger interactions with lipid membranes.^41^ However, we recently found that extensively S-alkylated ds DNA is structurally and thermally destabilized^42^ and that this is not a viable route to further increasing DLN stability.

Thus, we seek to explore alternative chemical modifications of DNA to interface with lipid bilayers that do not involve extensive charge neutralization or lead to aggregation. Previously, we found that DNA origamis that are coated with poly(ethylene) glycol (PEG) block copolymers are protected from enzymatic degradation.^43,44^ Moreover, PEG-modified origamis can even be quantitatively transferred across the phase boundary of a two-phase solution into organic solvents including chloroform, in which DNA is usually not soluble.^45^ The phase transfer takes place because the surface of the origamis is covered with PEG, which is an amphiphilic molecule that is even better soluble in chloroform than in water. As lipids are also commonly solubilized in chloroform, we hypothesized that the ethylene groups of the amphiphilic PEG molecules can provide an interface with the hydrophobic acyl chains of lipid membranes of a DLN. In this work, we covalently modified DNA circles with PEG, optimized the length and number of PEG modifications, incorporated a transmembrane peptide and analyzed the resulting complexes and interfaces with atomistic molecular dynamic simulations.

## Results and Discussion

### Oligonucleotide PEGylation

For this, we designed a 21-base oligonucleotide with up to three reactive amine modifications; one each at the 5’ and 3’ ends, and one at a modified thymidine (Figure 1A). The amino groups were reacted with PEG-37 NHS esters (molecular weight ∼2 kDa) through a nucleophilic substitution resulting in a stable amide bond. In separate experiments, we also conjugated shorter PEG-3- and PEG-13-NHS esters or an oligonucleotide with only one amine modification to investigate the respective effects of length and number of PEG chains in stable nanodisc formation (Figure S1). The PEG-oligonucleotide was purified by reverse-phase high performance liquid chromatography (RP-HPLC) (Figure S2) and characterized by high resolution liquid chromatography-mass spectrometry (LC-MS) (Figure S3). In denaturing polyacrylamide gel electrophoresis of the oligonucleotides (PAGE, Figure 1B), the addition of PEG reduces their electrophoretic mobility.

**Figure 1.**
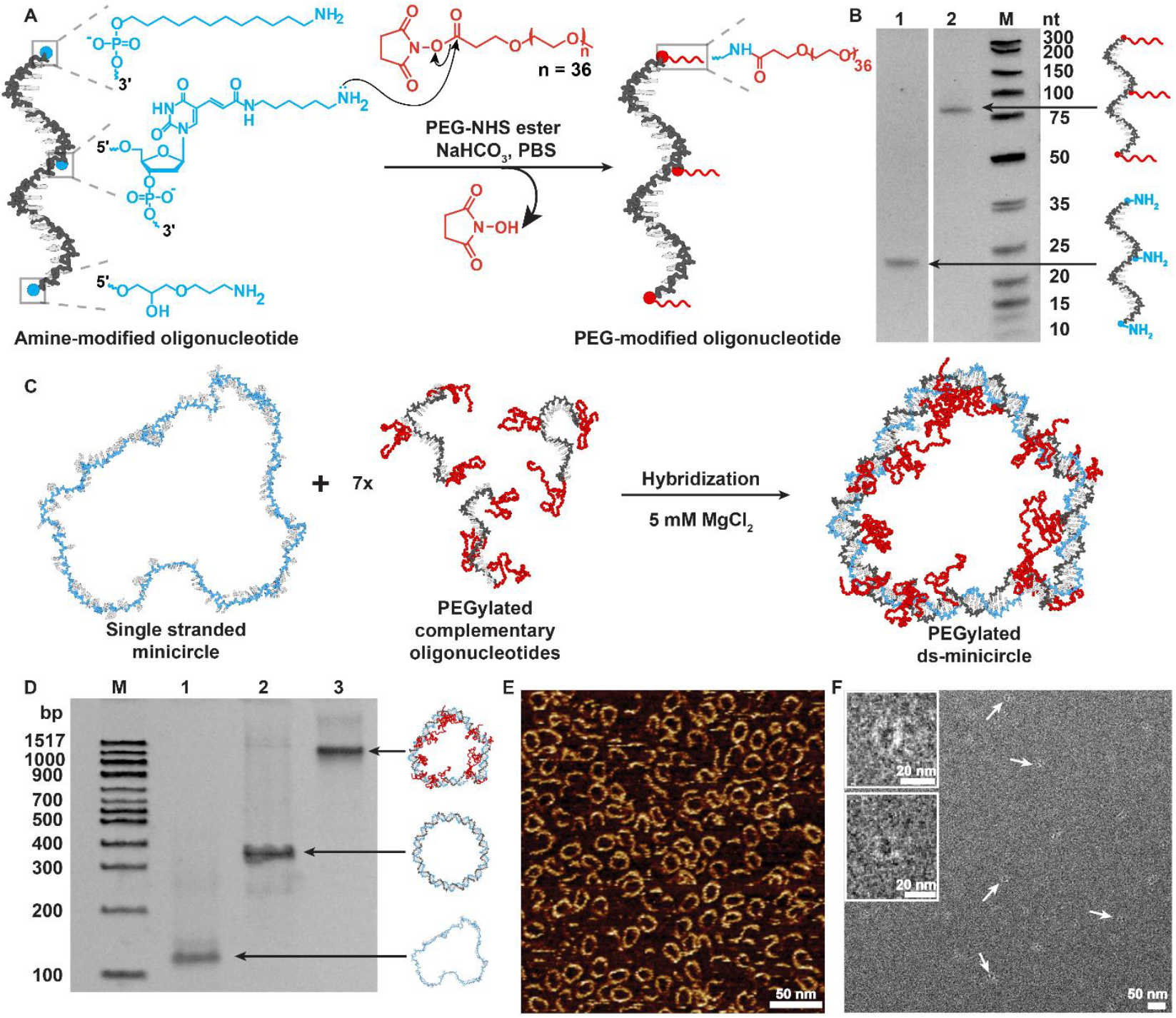
Synthesis and characterization of PEG-modified DNA minicircles. (A) A 21-mer oligonucleotide with three amine modifications was modified with PEG-NHS esters. (B) Denaturing PAGE of amine oligonucleotide, before and after reaction with PEG-37-NHS ester. (C) Hybridization of the circular single-stranded scaffold with seven identical complementary PEG-modified strands form a double-stranded (ds) circle. Shown are snapshots from MD simulations that illustrate the different flexibility of the polymers and ds DNA. (D) Native PAGE of the different circles. (M) marker, (1) ss DNA circle / scaffold, (2) ds circle formed with seven complementary unmodified oligonucleotides, (3) ds circles with seven complementary oligonucleotides, each modified with three PEG-37. AFM (E) and TEM (F) images of PEG-modified ds circles. Scale bar: 50 nm. Inset: zoomed-in image of ds minicircles.

### Formation PEGylated minicircle

We then hybridized the PEGylated oligonucleotides with a 147 nt (nucleotide) ss-minicircle scaffold that we prepared by splint ligation of a long strand (Figure 1C).^14^ The scaffold sequence contains seven repeats of an intrinsically bent A-tract motif^46^ that is complementary to the PEG-modified oligonucleotides. The ds-minicircles thus formed have an outer diameter of about 18 nm and an inner diameter of 14 nm (calculated using 0.34 nm/bp step) and are 2 nm thick. In a native polyacrylamide gel (PAGE), the modified ds circles form a sharp band with a lower electrophoretic mobility than ds minicircles without PEG modifications (Figure 1D). In atomic force microscopy (AFM) images (Figure 1E), and transmission electron microscopy (TEM) images (Figure 1F), the ds-PEG-minicircles appear intact and circular. The contrast of rings is higher in AFM than in TEM as DNA is not stained well in TEM. Moreover, the sample density is higher in AFM as rings adhere better to the positivized mica substrate than to TEM grids.

Note that this chemical coupling reaction does not neutralize charges in the DNA backbone as in S-alkylation of phosphorothioates^14,31^ that reduces their melting temperature (T_m_) and can destabilize hybridization.^42^ Despite the large added molecular mass, PEG-modification only lowered the melting temperature T_m_ with the ss minicircle by 4-5 °C (Figure S4).

### Synthesis of PEGylated DNA-lipid nanodiscs

To reconstitute a lipid bilayer within the PEGylated minicircle (Figure 2A), we mixed them with detergent-solubilized lipids and removed the detergents through a spin column. While attempts with zwitterionic detergents such as CHAPS or non-ionic detergents like Tween-20 did not result in DLNs, DLNs readily formed with positively charged detergents such as dodecyl trimethylammonium bromide (DTAB). This suggests that electrostatic interactions between mixed micelle and the negatively charged phosphate backbone of DNA are crucial for the formation of DLNs. The optimized lipid formulation contained 89 % DMPC (1,2-Dimyristoyl-sn-glycero-3-phosphocholine), which has a zwitterionic headgroup, 10 % DMTAP (1,2-dimyristoyl-3-trimethylammonium-propane) with a cationic head group and 1 % of the fluorescent lipid Topfluor-PC (TfPC). 10 % cationic lipid was found to be necessary for DLN formation (Figure S5) due to favorable electrostatic interactions with the phosphate backbone (Figure 3E-G). The detergent-solubilized lipids were mixed with dsDNA at a ratio of dsDNA: lipid = 1: 450, which was calculated to fill the area inside the DNA minicircles. A TEM image of unpurified DLNs after the detergent removal step (Figure 2B) reveals the successful synthesis of nanodiscs with a better imaging contrast than the empty ds-PEG-rings (Figure 1F). A control detergent removal experiment without PEGylated ds-minicircles was conducted, in which only liposomes formed as expected (Figure 2C).

**Figure 2.**
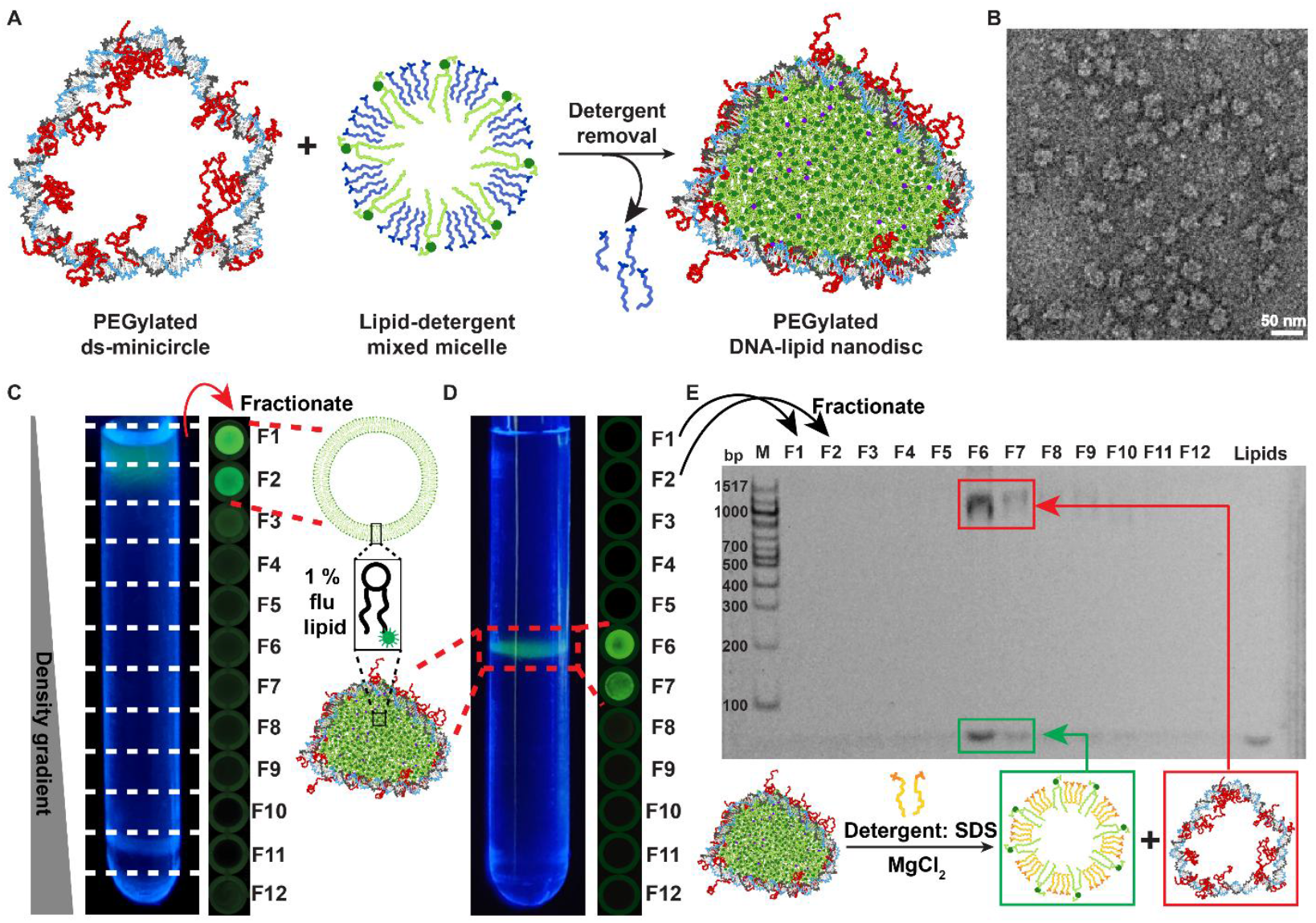
Synthesis and characterization of PEGylated DNA-lipid nanodiscs. (A) Lipid reconstitution in ds minicircles is achieved by adding lipid-detergent mixed micelles followed by detergent removal. (B) TEM image of ds minicircles with a lipid-filled interior. Scale bar: 50 nm. (C) Isopycnic ultracentrifugation of liposomes containing 1% fluorescent TfPC that were prepared by detergent removal without PEGylated DNA minicircles. The content of the ultracentrifugation tubes (left) was fractionated into the wells of a 96-well plate and imaged in a fluorescence scanner (right), indicating liposomes in the upper fractions (F1-F2). (D) Isopycnic centrifugation of DLNs showing lipids in F6-F7. (E) SDS PAGE analysis of the fractions taken from (D) shows confirms colocalization of PEGylated ds minicircle (red) and lipids (green) in fractions 6 and 7.

**Figure 3.**
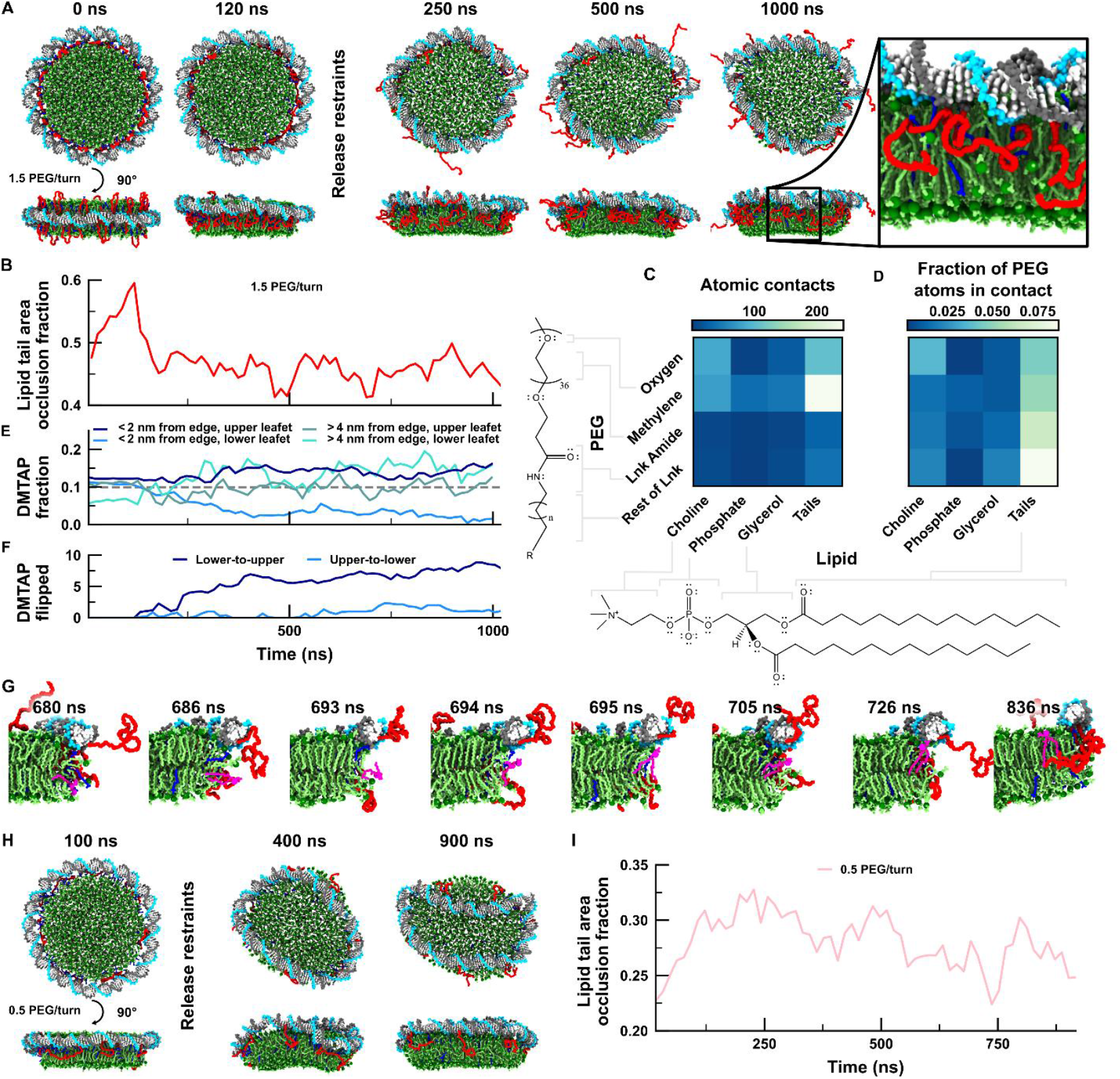
All-atom molecular dynamics simulations of PEGylated DLNs. (A) Snapshots of the restrained equilibration and unrestrained all-atom simulation trajectory of a DLN with 1.5 PEG-37 per DNA turn (21 total). (B) Fraction of lipid tail surface area occluded by PEG and DTAB in the DLN. The lipid tail area is calculated using a 2-Å probe radius, see Methods for details. (C,D) Contacts between lipid and PEG motifs, including linkers that tether PEG to the DNA. Both the average number of contacts (C) and fraction of possible pairs forming a contact (D) are depicted. (E) Ratio of DMTAP to all lipids, excluding DTAB, in four different regions of a DLN. The distance from the bilayer edge was computed as the lipid C2 atom distance from a 1-nm thick slice of the molecular surface representing the bilayer after projection into the plane of the bilayer, see Methods for details. The dashed gray line indicates the average DMTAP fraction in the system. A lipid was classified as being in the upper (lower) leaflet if its C2 atom was located above (below) the plane of the bilayer bisecting its center of mass. (F) Number of DMTAP lipids initially observed in the lower (upper) leaflet later observed in the upper (lower) leaflet. (G) Snapshots illustrating the flipping of a DMTAP lipid (pink). (H) Snapshots of an instable DLN with only seven instead of 21 PEG modifications (=0.5 PEG per helical turn). (I) Lipid tail area occlusion fraction as defined in panel B for the 0.5 PEG/turn system. All plotted timeseries were filtered with a 15 ns block average.

### Characterization of DLNs

Liposomes and DLNs were analyzed and purified through isopycnic ultracentrifugation in a density gradient made with iodixanol and buffer. Liposomes have a low density due to their large buffer-filled interior and float up to the top of the gradient (Figure 2C) whereas empty ds-PEG-rings stay at the bottom of the gradient (Figure S6) due to their high density (∼1.7 gcm^-3^). DLNs float up towards the middle, until their density matches that of the surrounding gradient. Under UV or blue light illumination, the Topfluor-PC in the lipid formulation fluoresces green. Additionally, the content of the ultracentrifugation tubes was fractionated into 96-well plates and imaged in a fluorescence scanner. While liposomes appeared in fractions 1-2, the lipids of DLNs were majorly found in fractions 6 and 7 (Figure 2C-D).

To prove the colocalization of DNA with the lipids in DLNs, we loaded aliquots of all fractions into a polyacrylamide gel that contained MgCl_2_ to stabilize dsDNA and the anionic detergent SDS that solubilizes the lipids into mixed micelles. DNA and lipids are therefore separated in the gel, but ds-rings remain intact. After gel electrophoresis, the gel was stained with the DNA stain SYBR gold.^47^ In this SDS gel, both ds-rings and lipids appeared in fractions 6 and 7 indicating stable complexes between the molecules (Figure 2E). A control lipid reconstitution experiment with ds-rings that were not PEG modified did not form DLNs (Figure S8), indicating that the PEG-37 modifications are necessary to hold a lipid bilayer. Staining the lipids with Nile red as an alternative to adding fluorescent lipids to the formulation failed due to strong non-specific interactions of the dye with iodixanol in the density gradient (Figure S7).

### Optimizing PEG content

Next, we reduced the number of PEG-37 modifications from three to one per oligonucleotide or seven PEG-37 per ds-ring to test how many PEG residues are necessary to hold the lipids in the ring, and no stable nanodiscs were observed for seven PEG modifications (Figure S9). Likewise, DLNs did not form with ds-circles containing shorter PEG-3 modifications and only inefficiently with PEG-13 chains (Figure S10), underlining the importance of longer PEG chains to stabilize the rim of the lipid bilayer.

Inspired by this result, we conjugated DNA to even longer PEG-5000 with an average molecular weight of 5 kDa. Nanodiscs only formed when the number of modifications per ring was reduced from 21 to seven (Figure S11). We hypothesize that the much longer and bulkier PEG-5000 chains occupy more of the center of the ring and act like an entropic brush that prevents the incorporation of lipids when DNA is extensively modified. These results indicate that the optimal functionalization is 21 PEG-37 chains per ds-minicircle or 1.5 PEG-37 modifications per helical turn.

### Further optimizations

Double-stranded DNA is much stiffer than single-stranded DNA, MSP or SMA scaffolds. We therefore incorporated seven 2-nt ss-gaps to create flexible joints to our minicircles. We hypothesized that these ss-gaps will cause the ring to be more compacted^48^ when empty, potentially facilitating nucleation of lipid mixed micelles during the lipid reconstitution. However, results were indistinguishable from experiments without gaps (Figure S12). We also identified that buffers used for the synthesis of other membrane mimetics (MSP, SMALP) are also suitable for the formation of the PEG-DLNs (Figure S13). However, high-salt buffers are detrimental to nanodisc formation (Figure S13) and thus we carried out DLN synthesis in buffers containing only 3 mM Mg^2+^, which is sufficient to stabilize the dsDNA without affecting the synthesis of the DLNs.

### MD Simulations of DLNs

Complementing the experimental characterization of DLNs, we constructed an all-atom system consisting of a patch of lipid bilayer surrounded by an idealized DNA ring functionalized with 1.5 PEG molecules per DNA turn to understand what molecular interactions take place at the interface between the lipid bilayer and the DNA. In addition, we partially neutralized the DNA-lipid interface with some cationic DTAB detergents to test if some DTAB might remain bound to the negatively charged phosphates at the DNA-lipid interface after the detergent removal step in experiments, potentially stabilizing the DNA-lipid interface. The system was submerged in ∼100 mM Na_2_SO_4_ and ∼6 mM MgCl_2_ electrolyte. During the first ∼120 ns of simulation, the non-hydrogen lipid atoms were harmonically restrained about their initial coordinates, and the DNA configuration was restrained via a stiff elastic network while the ions, DTAB, and PEG molecules were free to equilibrate. As restraints were gradually released, the entire DNA ring translated towards the upper leaflet of the bilayer (Figure 3A). After the restraints were released, the simulation was extended to one microsecond. The bilayer remained roughly circular whereas the DNA and PEG fluctuated extensively (Figure S15A; Movie 1). For comparison, an idealized bilayer patch was simulated without the DNA scaffold and also remained stable during the 1000-ns simulation (Movie 2). Note that in experiments such patches would fuse into lipid vesicles without additional stabilization of the edges.

Compared to the bare bilayer, the DNA-supported bilayer was marginally thinner and denser on average (Figure S15A, B). Moreover, the DNA, positioned on the upper leaflet, evidently induces a slight curvature (> 100 nm radius) away from the DNA (Figure S15C), but the differences in the average structure of the core (4 nm cylindrical region) of the nanodisc-supported bilayer and bare bilayer are negligible. However, the edges of the bilayers are distinct: the supported bilayer exhibited ∼30% fewer lipid headgroups along the edge than the bare bilayer (Figure S15D). The reduction of lipids along the edges indicates that the PEG polymers function as designed, and 45% of the surface area of the lipid tails is occluded by PEG, (Figure 3B). Moreover, the total lipid tail surface area in the DLN is reduced to ∼75% of that exposed in the bare bilayer (16,400 vs 21,500 Å^2^), despite their identical lipid compositions, indicating that the DNA scaffold reduces solvent-accessible crypts between headgroups. Most molecular interactions occur between lipid tails and the methylene in PEG (Figure 3C), which are abundant in both molecules and are the main factor stabilizing the DNA-PEG nanodiscs. Normalizing by the total number of atom pairs for each type of contact, the contacts between PEG oxygen and lipid choline are the most probable, followed by glycerol—glycerol interactions involving the linker regions (Figure 3D). Strikingly, the interactions with lipid phosphates are quite rare in both absolute and relative senses.

DLN synthesis was only successful using 10% of cationic lipids (DMTAP) suggesting an electrostatic interaction between DNA scaffold and the charged lipids in the bilayer. In simulations, we indeed observed a ∼50% enrichment of the lipid fraction of charged DMTAP near the DNA on the upper leaflet, and an even greater depletion of DMTAP in the same area on the opposing leaflet (Figure 3E), indicating migration of DMTAP towards the DNA. Towards the end of the simulation, DMTAP was enriched in the central region of the bilayer of the lower leaflet, which might be attributed to electrostatic interactions across the periodic boundary of the system. DMTAP lipids flipped almost exclusively from lower to upper leaflet (Figure 3F,G), with approximately five migrating during the trajectory against a background of ∼45 DMPC lipids flipping in each direction.

In contrast, the smaller cationic DTAB detergent molecules that were initially placed at the interface between the DNA and lipid bilayer did not show a strong affinity to the DNA (Figure S16) but quickly diffused into the surrounding water or into the lipid bilayer. This suggests that while DTAB facilitates the lipid reconstitution from mixed micelles in experiments, it is not an important factor for the stabilization of the interface between DNA and lipids once DLNs have formed.

Finally, we simulated the system with only seven instead of 21 PEG chains in the dsDNA circle which did not form DLNs in experiments. Likewise, the simulated DLN appeared less stable as the DNA slipped over the edge of the lipid bilayer (Figure 3H), although the PEG molecules remained associated with the lipid bilayer, with each PEG molecule occluding roughly twice as much lipid surface area per molecule as in the 1.5 PEG/turn system (Figure 3I). This further demonstrates the high affinity of PEG to the lipid edge, but the number of PEGs appears insufficient to stably hold the DNA ring in place. The bilayer thickness, curvature, and the number of lipids observed along the bilayer edge were similar to the unsupported bilayer (Figure S15).

### Incorporation of transmembrane peptide

Bilayer mimetics are often used for the biophysical characterization of membrane proteins^49^ or for drug delivery of hydrophobic molecules.^50,51^ As proof of principle for such future applications, we incorporated the hydrophobic transmembrane domain (TMD) of synaptobrevin, a protein that is found in the membrane of synaptic vesicles and is involved in the formation of the SNARE complex.^52^ For this, we modified the C-terminus of the TMD peptide with a biotin and bound it to a streptavidin-modified quantum dot (QD). The QD facilitates ultracentrifugation analysis due to its high density, its red fluorescence emission (λ_em_ = 655 nm) and produces high-contrast features in TEM images. All components including DTAB-solubilized lipids and peptides, PEG-DLNs, and streptavidin-QDs were mixed followed by detergent removal (Figure 4A). The complexes were characterized by isopycnic centrifugation as described above. The resulting QD-modified DLN produced a band with both green fluorescence from the Tf-PC lipid and red fluorescence from the QD and shifted from F6 to F7 due to the high density from the added QD (Figure 4B). TEM images of fraction 7 (Figure 4C) clearly show QDs bound to the DLNs with uniform sizes of 16±2 nm, confirming the integration of the TM peptide into the lipid bilayer. Without PEG modifications or the biotinylated TMD peptide, QDs did not co-localize with the dsDNA rings (Figure S14), proving the incorporation of the TMD peptide in the DLN.

**Figure 4.**
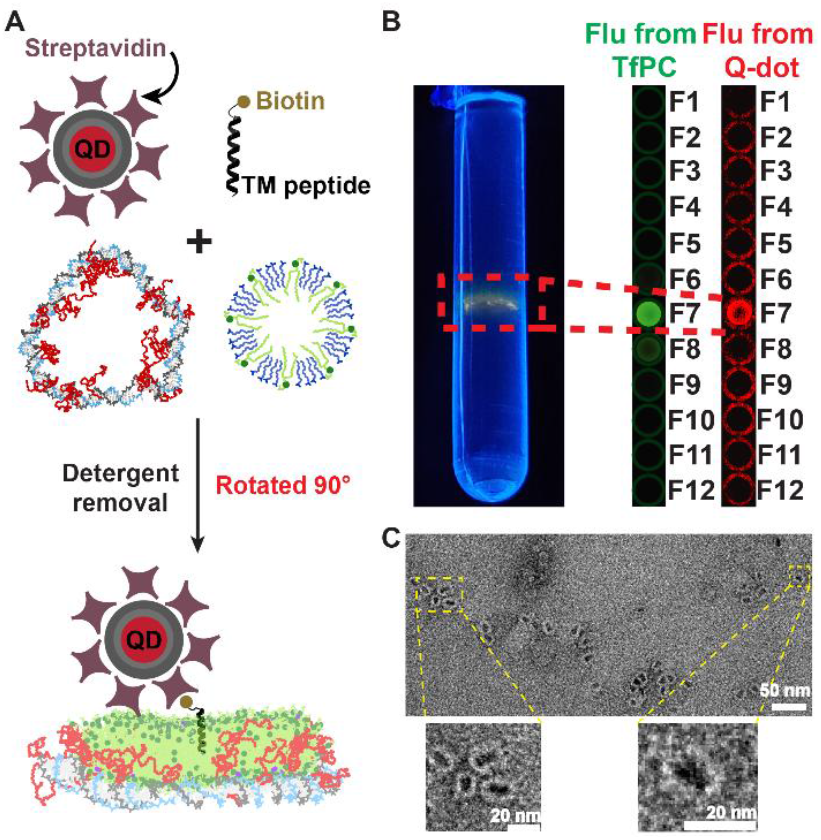
Incorporation of a transmembrane (TM) peptide into DLNs. (A) Incorporation of a biotin-modified transmembrane peptide in the lipid bilayer where biotin binds to a streptavidin-modified quantum dot (QD). (B) Isopycnic centrifugation of the DLN with the peptide and QD showing colocalized signals from fluorescent lipid and the streptavidin QDs indicating incorporation of the peptide in the DLN. (C) TEM image of DLN with a bound QD. Scale bar: 50 nm. Scale bar for zoomed-in images: 20 nm

## Conclusion

In this study, we developed a poly(ethylene glycol) modified DNA-based membrane mimetic to envelop lipid bilayers exploiting the favorable interactions between amphiphilic PEG chains and lipids. The conjugation of amphiphilic PEG chains of various lengths to multiple positions had little impact on the T_m_ of DNA to its template, but a minimum of three PEG 37 was necessary for DLN formation. Both reducing the length of PEG modifications and/or reducing the number of modifications significantly impacted the efficiency of lipid incorporation within the minicircles, while much longer PEG molecules inhibited incorporation of lipids. The optimized DNA circles contained 1.5 PEG 37 chains per helical turn and in ultracentrifugation experiments were quantitatively filled with lipids to form PEG-DLNs. Our atomistic MD simulations show qualitatively the same trends as experiments. They reveal that the PEG-minicircles stably hold lipids, but the DNA asymmetrically slipped towards the polar lipid head groups. The methylene groups of PEG chains interacted with the hydrophobic tails of the lipid bilayer, occluding a significant fraction of the solvent-exposed lipid tail surface area and mitigating the number of lipids that would roll over the edges of the bilayer. Additionally, the simulations reveal a flipping of positively charged lipids to the leaflet that is closest to the DNA circle and an enrichment of these positive lipids at the edge of the bilayer where they interact with the negatively charged phosphate backbone.

As a proof-of-concept, we incorporated the transmembrane domain of synaptobrevin that were bound to large streptavidin-modified QDs into our PEG-DLNs suggesting that large membrane proteins could also be incorporated in future structural studies. We envision that PEG-DLNs enable applications in structural biology, biophysics, and biomedicine due to the modularity of the design and the possibility to extend the design with additional, site-specific functionalizations.

## Associated Content

Supporting information contains additional experimental materials and methods, computational methods, electrophoresis data, HPLC chromatograms, MS data, Tm data, ultracentrifugation results and additional simulation analyses, movies of the simulation trajectories and descriptions of the movies.

## AUTHOR INFORMATION

### Author Contributions

T.L.S. and S.C. conceived the study; S.C. and S.K. performed experiments; C.M. performed all-atom molecular dynamics simulations; J.K. and D.P.N.G. synthesized the modified transmembrane peptide; all authors analyzed and visualized the data; T.L.S. and A.A. acquired funding; T.L.S. supervised the study; S.C., C.M. and T.L.S. wrote the initial draft; all authors revised and expanded the manuscript.

### Notes

The authors declare no competing financial interest

## Acknowledgments

We thank Martin Held and Dr. Alexander Heckel for MS characterization of the PEG oligonucleotide conjugates. We thank Dr. Edgar Kooijman for helpful discussions. We thank Praneetha Sundar Prakash for experiments involving DLN synthesis with different lipid mixtures.

## References

(1) Harayama, T.; Riezman, H. Understanding the Diversity of Membrane Lipid Composition. Nat Rev Mol Cell Biol 2018, 19 (5), 281–296. 10.1038/nrm.2017.138.

(2) Rigaud, J.-L.; Lévy, D. Reconstitution of Membrane Proteins into Liposomes. In Methods in Enzymology; Elsevier, 2003; Vol. 372, pp 65–86. 10.1016/S0076-6879(03)72004-7.

(3) Smirnova, I. A.; Ädelroth, P.; Brzezinski, P. Extraction and Liposome Reconstitution of Membrane Proteins with Their Native Lipids without the Use of Detergents. Scientific reports 2018, 8 (1), 14950. 10.1038/s41598-018-33208-1.

(4) Dürr, U. H. N.; Gildenberg, M.; Ramamoorthy, A. The Magic of Bicelles Lights Up Membrane Protein Structure. Chem. Rev. 2012, 112 (11), 6054–6074. 10.1021/cr300061w.

(5) Borch, J.; Hamann, T. The Nanodisc: A Novel Tool for Membrane Protein Studies. Biol Chem 2009, 390 (8), 805–814. 10.1515/BC.2009.091.

(6) Denisov, I. G.; Sligar, S. G. Nanodiscs in Membrane Biochemistry and Biophysics. Chem. Rev. 2017, 117 (6), 4669–4713. 10.1021/acs.chemrev.6b00690.

(7) Sligar, S. G.; Denisov, I. G. Nanodiscs: A Toolkit for Membrane Protein Science. Protein Science 2021, 30 (2), 297–315. 10.1002/pro.3994.

(8) Denisov, I. G.; Sligar, S. G. Nanodiscs for Structural and Functional Studies of Membrane Proteins. Nat Struct Mol Biol 2016, 23 (6), 481–486. 10.1038/nsmb.3195.

(9) Bayburt, T. H.; Sligar, S. G. Membrane Protein Assembly into Nanodiscs. FEBS Letters 2010, 584 (9), 1721–1727. 10.1016/j.febslet.2009.10.024.

(10) Kariyazono, H.; Nadai, R.; Miyajima, R.; Takechi-Haraya, Y.; Baba, T.; Shigenaga, A.; Okuhira, K.; Otaka, A.; Saito, H. Formation of Stable Nanodiscs by Bihelical Apolipoprotein A-I Mimetic Peptide. Journal of Peptide Science 2016, 22 (2), 116–122. 10.1002/psc.2847.

(11) Ikeda, K.; Nakano, M. Self-Reproduction of Nanoparticles through Synergistic Self-Assembly. Langmuir 2015, 31 (1), 17–21. 10.1021/la503491p.

(12) Mishra, V. K.; Anantharamaiah, G. M.; Segrest, J. P.; Palgunachari, M. N.; Chaddha, M.; Sham, S. W. S.; Krishna, N. R. Association of a Model Class A (Apolipoprotein) Amphipathic α Helical Peptide with Lipid: HIGH RESOLUTION NMR STUDIES OF PEPTIDE·LIPID DISCOIDAL COMPLEXES*. Journal of Biological Chemistry 2006, 281 (10), 6511–6519. 10.1074/jbc.M511475200.

(13) Postis, V.; Rawson, S.; Mitchell, J. K.; Lee, S. C.; Parslow, R. A.; Dafforn, T. R.; Baldwin, S. A.; Muench, S. P. The Use of SMALPs as a Novel Membrane Protein Scaffold for Structure Study by Negative Stain Electron Microscopy. Biochimica et Biophysica Acta (BBA) - Biomembranes 2015, 1848 (2), 496–501. 10.1016/j.bbamem.2014.10.018.

(14) Iric, K.; Subramanian, M.; Oertel, J.; Agarwal, N. P.; Matthies, M.; Periole, X.; Sakmar, T. P.; Huber, T.; Fahmy, K.; Schmidt, T. L. DNA-Encircled Lipid Bilayers. Nanoscale 2018, 10 (39), 18463–18467. 10.1039/C8NR06505E.

(15) Bayburt, T. H.; Grinkova, Y. V.; Sligar, S. G. Self-Assembly of Discoidal Phospholipid Bilayer Nanoparticles with Membrane Scaffold Proteins. Nano Lett. 2002, 2 (8), 853–856. 10.1021/nl025623k.

(16) Marconnet, A.; Michon, B.; Le Bon, C.; Giusti, F.; Tribet, C.; Zoonens, M. Solubilization and Stabilization of Membrane Proteins by Cycloalkane-Modified Amphiphilic Polymers. Biomacromolecules 2020, 21 (8), 3459–3467. 10.1021/acs.biomac.0c00929.

(17) Smith, A. A. A.; Autzen, H. E.; Faust, B.; Mann, J. L.; Muir, B. W.; Howard, S.; Postma, A.; Spakowitz, A. J.; Cheng, Y.; Appel, E. A. Lipid Nanodiscs via Ordered Copolymers. Chem 2020, 6 (10), 2782–2795. 10.1016/j.chempr.2020.08.004.

(18) Oluwole, A. O.; Danielczak, B.; Meister, A.; Babalola, J. O.; Vargas, C.; Keller, S. Solubilization of Membrane Proteins into Functional Lipid-Bilayer Nanodiscs Using a Diisobutylene/Maleic Acid Copolymer. Angew. Chem. Int. Ed. 2017, 56 (7), 1919–1924. 10.1002/anie.201610778.

(19) Zhan, P.; Peil, A.; Jiang, Q.; Wang, D.; Mousavi, S.; Xiong, Q.; Shen, Q.; Shang, Y.; Ding, B.; Lin, C.; Ke, Y.; Liu, N. Recent Advances in DNA Origami-Engineered Nanomaterials and Applications. Chem. Rev. 2023. 10.1021/acs.chemrev.3c00028.

(20) Madsen, M.; Gothelf, K. V. Chemistries for DNA Nanotechnology. Chem. Rev. 2019, 119 (10), 6384–6458. 10.1021/acs.chemrev.8b00570.

(21) Kocabey, S.; Kempter, S.; List, J.; Xing, Y.; Bae, W.; Schiffels, D.; Shih, W. M.; Simmel, F. C.; Liedl, T. Membrane-Assisted Growth of DNA Origami Nanostructure Arrays. ACS Nano 2015, 9 (4), 3530–3539. 10.1021/acsnano.5b00161.

(22) List, J.; Weber, M.; Simmel, F. C. Hydrophobic Actuation of a DNA Origami Bilayer Structure. Angew. Chem. Int. Ed. 2014, 53 (16), 4236–4239. 10.1002/anie.201310259.

(23) Sproat, B. S.; Rupp, T.; Menhardt, N.; Keane, D.; Beijer, B. Fast and Simple Purification of Chemically Modified Hammerhead Ribozymes Using a Lipophilic Capture Tag. Nucleic Acids Res 1999, 27 (8), 1950–1955. 10.1093/nar/27.8.1950.

(24) Perrault, S. D.; Shih, W. M. Virus-Inspired Membrane Encapsulation of DNA Nanostructures To Achieve In Vivo Stability. ACS Nano 2014, 8 (5), 5132–5140. 10.1021/nn5011914.

(25) Yang, Y.; Wang, J.; Shigematsu, H.; Xu, W.; Shih, W. M.; Rothman, J. E.; Lin, C. Self-Assembly of Size-Controlled Liposomes on DNA Nanotemplates. Nat. Chem. 2016, 8 (5), 476–483. 10.1038/nchem.2472.

(26) Börjesson, K.; Tumpane, J.; Ljungdahl, T.; Wilhelmsson, L. M.; Nordén, B.; Brown, T.; Mårtensson, J.; Albinsson, B. Membrane-Anchored DNA Assembly for Energy and Electron Transfer. J. Am. Chem. Soc. 2009, 131 (8), 2831–2839. 10.1021/ja8038294.

(27) Börjesson, K.; Lundberg, E. P.; Woller, J. G.; Nordén, B.; Albinsson, B. Soft-Surface DNA Nanotechnology: DNA Constructs Anchored and Aligned to Lipid Membrane. Angew. Chem. Int. Ed. 2011, 50 (36), 8312–8315. 10.1002/anie.201103338.

(28) Burns, J. R.; Göpfrich, K.; Wood, J. W.; Thacker, V. V.; Stulz, E.; Keyser, U. F.; Howorka, S. Lipid-Bilayer-Spanning DNA Nanopores with a Bifunctional Porphyrin Anchor. Angew. Chem. Int. Ed. 2013, 52 (46), 12069. 10.1002/anie.201305765.

(29) Vyborna, Y.; Vybornyi, M.; Rudnev, A. V.; Häner, R. DNA-Grafted Supramolecular Polymers: Helical Ribbon Structures Formed by Self-Assembly of Pyrene–DNA Chimeric Oligomers. Angew. Chem. Int. Ed. 2015, 54 (27), 7934–7938. 10.1002/anie.201502066.

(30) Burns, J. R.; Stulz, E.; Howorka, S. Self-Assembled DNA Nanopores That Span Lipid Bilayers. Nano Lett. 2013, 13 (6), 2351–2356. 10.1021/nl304147f.

(31) Burns, J. R.; Al-Juffali, N.; Janes, S. M.; Howorka, S. Membrane-Spanning DNA Nanopores with Cytotoxic Effect. Angew. Chem. Int. Ed. 2014, 53 (46), 12466–12470. 10.1002/anie.201405719.

(32) Zhao, Z.; Zhang, M.; Hogle, J. M.; Shih, W. M.; Wagner, G.; Nasr, M. L. DNA-Corralled Nanodiscs for the Structural and Functional Characterization of Membrane Proteins and Viral Entry. Journal of the American Chemical Society 2018, 140 (34). 10.1021/jacs.8b04638.

(33) Czogalla, A.; Kauert, D. J.; Franquelim, H. G.; Uzunova, V.; Zhang, Y.; Seidel, R.; Schwille, P. Amphipathic DNA Origami Nanoparticles to Scaffold and Deform Lipid Membrane Vesicles. Angew. Chem. Int. Ed. 2015, 54 (22), 6501–6505. 10.1002/anie.201501173.

(34) Zhang, Z.; Yang, Y.; Pincet, F.; Llaguno, M. C.; Lin, C. Placing and Shaping Liposomes with Reconfigurable DNA Nanocages. Nat. Chem. 2017, 9 (7), 653–659. 10.1038/nchem.2802.

(35) Franquelim, H. G.; Khmelinskaia, A.; Sobczak, J.-P.; Dietz, H.; Schwille, P. Membrane Sculpting by Curved DNA Origami Scaffolds. Nat. Commun. 2018, 9 (1), 811.

(36) Aissaoui, N.; Mills, A.; Lai-Kee-Him, J.; Triomphe, N.; Cece, Q.; Doucet, C.; Bonhoure, A.; Vidal, M.; Ke, Y.; Bellot, G. Free-Standing DNA Origami Superlattice to Facilitate Cryo-EM Visualization of Membrane Vesicles. J. Am. Chem. Soc. 2024, 146 (19), 12925–12932. 10.1021/jacs.3c07328.

(37) Zhang, Z.; Feng, Z.; Zhao, X.; Yu, Z.; Chapman, E. R. Programmable Liposome Organization via DNA Origami Templates. J. Am. Chem. Soc. 2025, 147 (28), 24548–24554. 10.1021/jacs.5c05196.

(38) Xu, W.; Wang, J.; Rothman, J. E.; Pincet, F. Accelerating SNARE-Mediated Membrane Fusion by DNA–Lipid Tethers. Angew. Chem. Int. Ed. 2015, 54 (48), 14388–14392. 10.1002/anie.201506844.

(39) Stengel, G.; Zahn, R.; Höök, F. DNA-Induced Programmable Fusion of Phospholipid Vesicles. J. Am. Chem. Soc. 2007, 129 (31), 9584–9585. 10.1021/ja073200k.

(40) Pinner, M. T.; Dietz, H. Programmable DNA Shell Scaffolds for Directional Membrane Budding. Nat Commun 2025, 16 (1), 8972. 10.1038/s41467-025-64298-x.

(41) Maingi, V.; Rothemund, P. W. K. Properties of DNA- and Protein-Scaffolded Lipid Nanodiscs. ACS Nano 2021, 15 (1), 751–764. 10.1021/acsnano.0c07128.

(42) Chandrasekhar, S.; Bricker, R.; Fadaei, F.; Schmidt, T.-L. S-Alkyl-Phosphorothioate Modifications Reduce Thermal and Structural Stability of DNA Duplexes. bioRxiv September 30, 2025, p 2025.09.30.679560. 10.1101/2025.09.30.679560.

(43) Agarwal, N. P.; Matthies, M.; Gür, F. N.; Osada, K.; Schmidt, T. L. Block Copolymer Micellization as a Protection Strategy for DNA Origami. Angew. Chem. Int. Ed. 2017, 56 (20), 5460–5464. 10.1002/anie.201608873.

(44) Ponnuswamy, N.; Bastings, M. M. C.; Nathwani, B.; Ryu, J. H.; Chou, L. Y. T.; Vinther, M.; Li, W. A.; Anastassacos, F. M.; Mooney, D. J.; Shih, W. M. Oligolysine-Based Coating Protects DNA Nanostructures from Low-Salt Denaturation and Nuclease Degradation. Nature Communications 2017, 8, 15654. 10.1038/ncomms15654.

(45) Agarwal, N. P.; Chandrasekhar, S.; Prakash, P. S.; Joffroy, K.; Schmidt, T. L. Block Copolymer Micellization of DNA Origami Promotes Solubility in Organic Solvents. Langmuir 2022, 38 (38), 11650–11657. 10.1021/acs.langmuir.2c01508.

(46) MacDonald, D.; Herbert, K.; Zhang, X.; Polgruto, T.; Lu, P. Solution Structure of an A-Tract DNA Bend. J. Mol. Biol. 2001, 306 (5), 1081–1098. 10.1006/jmbi.2001.4447.

(47) Kolbeck, P. J.; Vanderlinden, W.; Gemmecker, G.; Gebhardt, C.; Lehmann, M.; Lak, A.; Nicolaus, T.; Cordes, T.; Lipfert, J. Molecular Structure, DNA Binding Mode, Photophysical Properties and Recommendations for Use of SYBR Gold. Nucleic Acids Research 2021, 49 (9), 5143–5158. 10.1093/nar/gkab265.

(48) Agarwal, N. P.; Matthies, M.; Joffroy, B.; Schmidt, T. L. Structural Transformation of Wireframe DNA Origami via DNA Polymerase Assisted Gap-Filling. ACS Nano 2018, 12 (3), 2546–2553. 10.1021/acsnano.7b08345.

(49) Majeed, S.; Ahmad, A. B.; Sehar, U.; Georgieva, E. R. Lipid Membrane Mimetics in Functional and Structural Studies of Integral Membrane Proteins. Membranes 2021, 11 (9), 685. 10.3390/membranes11090685.

(50) Peetla, C.; Stine, A.; Labhasetwar, V. Biophysical Interactions with Model Lipid Membranes: Applications in Drug Discovery and Drug Delivery. Mol. Pharmaceutics 2009, 6 (5), 1264–1276. 10.1021/mp9000662.

(51) Sheikhpour, M.; Barani, L.; Kasaeian, A. Biomimetics in Drug Delivery Systems: A Critical Review. Journal of Controlled Release 2017, 253, 97–109. 10.1016/j.jconrel.2017.03.026.

(52) Zhang, Z.; Chapman, E. R. Programmable Nanodisc Patterning by DNA Origami. Nano Lett. 2020, 20 (8), 6032–6037. 10.1021/acs.nanolett.0c02048.

