## Supporting information for "DNA-Lipid Nanodiscs with a Polyethylene Glycol Interface"

#### Experimental Methods

##### 1. Experimental Section

###### 1.1. Materials

All amine modified DNA oligonucleotides were purchased from Integrated DNA technologies in the desalted form and were suspended in deionized water to form a 1 mM solution. They were purified in-house with reverse-phase high performance liquid chromatography (RP-HPLC) from Agilent using a Restek Viva 5 µm C4 column (cat. no. 9512525). The oligonucleotides were then dried using a vacuum centrifuge (Eppendorf) followed by resuspension in water to form a 1 mM solution. The yield of the purified oligonucleotides varied between 50-70%. PEG-3-NHS (cat. no. BP-20980), PEG-13-NHS (cat. no. BP-22584) and PEG-37-NHS esters (cat. no. BP-22248) were obtained from Broadpharm. DMF (cat. no. 043465) and sodium bicarbonate (cat. no. 470302-440) were obtained from Alfa Aesar and Ward's science respectively. 10X PBS (cat. no. J75889-AE) was purchased from Thermo Scientific. 1,2-dimyristoyl-sn-glycero-3-phosphocholine (DMPC, cat. no. 850345),1,2-dimyristoyl-3-trimethylammonium-propane (chloride salt) (DMTAP, cat. no. 890860) and 1-palmitoyl-2-(dipyrrometheneboron difluoride)undecanoyl-sn-glycero-3-phosphocholine (Topfluor (Tf) PC, cat. no. 810281) were purchased in the powder form from Avanti Research. Dodecyltrimethyl ammonium bromide (DTAB, cat. no. D5047) was purchased from Sigma Aldrich. Optiprep density gradient medium (cat. no. AB286850) was obtained from Abcam. 96-well non-treated, sterilized plates were purchased from VWR (cat. no. 10861-562). QD 655 streptavidin conjugate (cat. no. Q10123MP) was obtained from Thermo Fisher Scientific. Chloroform (cat. no. CX1054-6) and methanol (cat. no. MX0488-1) were purchased from Millipore Sigma. Zymo-Spin IICR columns (cat. no. C1078), Oligo binding buffer (cat. no. D4060-1-40) and DNA wash buffer (cat. no. D4003-2-48) were purchased from Zymo research.

20% denaturing polyacrylamide gels were hand cast using 8 M Urea (VWR, cat. no. 0568), 10X TBE prepared in-house, 40% acrylamide-bisacrylamide solution (19:1) from Thermo Scientific (cat. no.

J60909.K2), 10% ammonium persulphate from VWR (cat. no. M133) and TEMED from Bio Rad (cat. no. 161-0801). GeneRuler Ultra Low Range DNA ladder was purchased from Thermo Fisher Scientific (cat. no. SM1211).

5% native gels with 5 mM $MgCl_2$ were hand cast using 40% acrylamide-bisacrylamide solution (29:1) from VWR (cat. no. 0311) and $MgCl_2$ hexahydrate (VWR, cat. no. BDH9244). All gels were run in 1X TBE (100 mM TRIS, 100 mM Boric Acid, 2 mM EDTA, pH 8.3). Tris(hydroxymethyl)amino methane was purchased from Sigma Aldrich (cat. no. 252859), Boric acid (cat. no. BDH9222) and disodium salt of EDTA were purchased from VWR (cat. no. BDH4616). SYBR gold (cat. no. S11494) and SYBR green (cat. no. S7563) DNA gel stains were purchased from Invitrogen by Thermo Fisher Scientific. HEPES (cat. no. J848) was purchased from VWR, $MgSO_4.7H_2O$ (cat. no. M2773) and $Na_2SO_4$ (cat. no. 71959) were purchased from Sigma Aldrich. 6X native DNA loading dye (cat. no. B7024S) and 100 bp DNA ladder (cat. no. N3231) were purchased from New England Biolabs. Uranyl formate hydrate was purchased from Electron Microscopy Sciences (cat. no. 22450).

##### 1.2. Experimental Methods

**PEGylation of amine-DNA.** 5 µL of a 1 mM solution of HPLC-purified 3-amine oligonucleotides were mixed with 1.625 µL of 10X PBS pH 7.4 and 1.625 µL of 1 M $NaHCO_3$ in a DNA-low bind microcentrifuge tube. This step causes deprotonation of the amine group making it suitable for nucleophilic attack of the lone pair at the carbonyl group of the NHS ester. To this solution, 8.04 µL of PEG-37 NHS ester (30-fold molar excess per amine) dissolved in anhydrous, amine-free DMF to a concentration of 100 mg/mL were added and mixed well. This reaction mixture with a total volume of ~16.3 µL was incubated in a thermomixer at 25 °C, 1000 rpm for 12 h.

Post-reaction, the sample was diluted with water to 100 µL and purified by RP-HPLC. The respective peaks were collected, and solvent was evaporated in a vacuum centrifuge and pellet was redissolved to make a 100 µM solution. The yields of the purified product were measured by UV absorbance and were about 80-90 %.

Conjugation of the 1-amine oligonucleotide to PEG-37-NHS was carried out in a similar manner but the volume of NHS ester was proportionately lowered to 2.68 µL. Conjugation of 1- and 3-amine oligonucleotides to PEG-3 and PEG-13 NHS esters were also performed simultaneously according to the protocol described above.

Sequences of amine oligonucleotides using the modifier codes from IDT DNA.

1-amine – ACACTTTTTTCACACTTTTTTC/3AmMO/

3-amine – /5AmMC12/ACACTTTTT/iAmMC6T/CACACTTTTTC/3AmMO/

**Denaturing polyacrylamide gel electrophoresis.** 20% denaturing urea polyacrylamide gels were cast using 5 mL of 40 % acrylamide-bisacrylamide (19:1), 1 mL of 10X TBE, 4.8 g of urea, 100 µL of 10 % APS, 10 µL of TEMED and water to make up the total volume to 10 mL. Gels were polymerized for 30-60 min before samples were loaded. 7.5 µL of 20 pmol of starting amine-DNA and the PEGylated product were each mixed with 2.5 µL of 4X denaturing loading dye (100% formamide, 10 mM NaOH, 0.01 % bromophenol blue, 0.02 % xylene cyanol) and loaded in the sample wells. 1 µL of 0.01 µg/mL of the GeneRuler ultralow range DNA ladder was also mixed with the denaturing loading dye and applied to sample wells as a reference. Electrophoresis was performed at 200 V for 65 min at 65 °C.

**Formation of single-stranded circular scaffold and double-stranded minicircles.** Single-stranded circles were synthesized according to a previously published protocol.[1] Briefly, a 147-nt linear oligonucleotide was phosphorylated at the 5' end using T4 phosphonucleotide kinase and the ends were splint ligated using T4 DNA ligase. Uncircularized strands and the splints were removed by digestion with Exonuclease I and III. Pure circular product was purified by affinity columns (Zymo research) followed by yield estimation using absorbance measurements.

The sequence of single-stranded 147-mer DNA was - /5Phos/$(TGTGAAAAAGTGTGAAAAAG)_7$

Double-stranded (ds) minicircles were formed by annealing 500 pmol of the circular scaffold with 12.5 nmol of the complementary unmodified or PEGylated oligonucleotides in 5 mM $MgCl_2$ by heating to 80 °C and slowly cooling them to 25 °C over an hour. The ds-circles were purified with 50 kDa molecular weight cut-off (MWCO) filters in a buffer containing 50 mM HEPES, 3 mM $MgSO_4$ and 100 mM $Na_2SO_4$. The sample was washed 5-7 times with 500 µL of buffer and spun at 10000 rcf for 5 min. Finally, the filter was inverted and the sample was retrieved by spinning at 10000 rcf for 3 min. Yields were determined by measuring absorbance at 260 nm and the ds-circles were analyzed by native polyacrylamide gel electrophoresis (PAGE) or AFM.

**Native polyacrylamide gel electrophoresis.** 5 % native polyacrylamide gels were cast using 1.25 mL of 40 % acrylamide-bisacrylamide (29:1), 1 mL of 10X TBE, 250 µL of 200 mM $MgCl_2$, 100 µL of 10 % APS, 10 µL of TEMED and 7.39 mL of water to make up the total volume to 10 mL. Native-SDS gels consisted of an additional 100 µL of 5 % SDS in them. Gels were allowed to polymerize for 30-60 min before samples were loaded. 10 µL of 0.5 pmol of single-stranded circular scaffold were mixed with 2 µL of commercial 6X native loading dye (2.5 % Ficoll-400, 10 mM EDTA, 3.3 mM Tris-HCl, 0.08 % SDS, 0.02 % Dye 1, 0.001 % Dye 2, pH 8.0 at 25 °C) was loaded in the sample wells. For native-SDS gels, 10 µL of fractions after ultracentrifugation were mixed with 1.1 µL of 10X native-SDS dye prepared in-house

(100 mM Tris-HCl, 50 % glycerol, 0.5 % SDS, pH 8.0 at 25 °C) and loaded. Since SYBR gold stains dsDNA better than ss DNA, only 0.25 pmol of ds-rings were loaded into gels. 1 µL of 50 µg/mL of the 100 bp DNA ladder was also mixed with the native/native-SDS loading dye and applied to sample wells as a reference. Electrophoresis was performed at 100 V for 65 min at 4 °C in a running buffer containing 1X TBE, 5 mM $MgCl_2$ for native gels and an additional 0.05 % SDS for native-SDS gels.

**Denaturing and native PAGE staining and imaging.** When electrophoresis was complete, the gel was stained in 1X SYBR gold in 1X TBE supplemented with 5% ethanol to prevent the stain from sticking to the plastic staining tray. Gels were then imaged on a GE Typhoon FLE 9500 gel scanner using a 473 nm excitation laser and 510 nm long pass emission filter. The photomultiplier tube gain was set to 500 and the pixel size was 50 µm. Images were analyzed by the Fiji ImageJ image analysis software.

**Atomic Force Microscopy.** 20 µL of a 0.01% polyornithine (Sigma Aldrich, cat. no. P3655) solution were added onto freshly cleaved mica and incubated for 2 min. The mica surface was then washed with ~10 mL of Milli-Q and dried with a stream of $N_2$ gas. 10 µL of unmodified or PEGylated ds-rings (20 nM) were deposited onto the mica (Ted Pella, 9.9 mm diameter) and incubated for 3–5 min. The solution was wicked away and the surface was washed twice with 1 mL of Milli-Q water after which 60 µL of water were added on to the sample surface. 20 µL of water were also added onto the ScanAsyst Fluid+ tip (Bruker, cat. no. p-3728) and imaging was performed in liquid using the peak force tapping mode in a highspeed atomic force microscope (Bruker, Dimension Fastscan bio). Images were processed with the Nanoscope analysis software.

**Transmission Electron Microscopy.** 400-mesh carbon-only grids (Electron Microscopy Sciences, cat no. CF400-Cu-UL) were glow-discharged for 30 s at 15 mA negative polarity using a PELCO easiGlow glow discharge system to render the surface hydrophilic and facilitate sample deposition in the subsequent steps. For optimum results, glow-discharged grids were used within 15 min of treatment. 5 µL of the purified ds-PEG-minicircles or PEG-DLNs or QD-peptide-PEG-DLNs were applied to treated grids and incubated for 5 min. Meanwhile, a fresh 1% uranyl formate solution in water was prepared. To this, 1.25 µL of freshly prepared 1 M NaOH were added, mixed well for 5 min in a thermomixer at 750 rpm, and centrifuged for 30 s using a tabletop microcentrifuge. We observed that preparation of fresh uranyl stain showed dramatic improvement in results compared to uranyl stains that were stored at −20 °C for extended periods of time. After the sample was incubated for 5 min, the excess sample was wicked away, and 10 µL of freshly activated uranyl formate were applied to the grid and incubated for 30 s before the excess was wicked away. The grids were then air-dried and imaged using a Tecnai F20 transmission electron microscope operated at 200 kV. Images were analyzed with the Fiji ImageJ analysis software.

**$T_m$ analysis.** A 21-mer DNA which is the exact complement of the PEGylated oligonucleotide was designed (sequence: GAAAAAGTGTGAAAAAAGTGT). 1 µL of the 21-mer DNA (200 µM) was mixed with 1 µL (5-fold molar excess) of the complementary unmodified or PEGylated oligonucleotide (1 mM) in 1X TE buffer with 5 mM $MgCl_2$. To this 3 µL of 10X SYBR green (such that final concentration was 1X) were added to record real-time change in fluorescence upon DNA hybridization. The total reaction volume was 30 µL. Additionally, samples were also layered with oil to prevent evaporation. Samples were placed in a real-time magnetic induction cycler (Biomolecular Systems) and heated to 85 °C for 2 min through magnetic induction followed by cooling to 40 °C over an hour using fan-forced air. Fluorescence from the DNA-binding dye increased as the temperature decreased due to formation of dsDNA and was recorded with high-sensitivity photodiodes for each dedicated excitation/emission channel. The micPCR software was used to take the first derivative of the change in fluorescence with respect to temperature (dF/dT) to obtain the $T_m$ of the duplex.

**Lipid film preparation.** Powder stocks of DMPC, DMTAP and TfPC were dissolved in 2:1 chloroform: methanol to make a stock concentration of 25 mg/mL for DMPC, DMTAP and 1 mg/mL for TfPC respectively. Stock solutions of lipids were used immediately and are not recommended for long-term storage. 12.06 µL of DMPC, 1.18 µL of DMTAP and 5 µL of TfPC in the ratio of 89: 10: 1 respectively, were mixed in a clean glass test tube. The solvent was evaporated using a stream of $N_2$ gas while the tube was placed in hot water to ensure all lipids remained above their phase transition temperature as they dry. The dried lipid films were placed in a vacuum desiccator overnight. Finally, the tubes were purged with $N_2$, sealed and stored at -20 °C until further use.

**Nanodisc preparation.** Lipids were dissolved to a final concentration of 2 mM in 1X Buffer H (50 mM HEPES, 3 mM $MgSO_4$ and 100 mM $Na_2SO_4$) with 60 mM DTAB and sonicated for 10 min to form mixed micelles. 8.53 µL (25 pmol) of DNA-PEG-minicircles in 1X buffer H were mixed with 5.62 µL (11.25 µmol) of lipids at a molar ratio of 1: 450 (DNA: lipid). 13.23 µL of 100 mM DTAB were added to adjust the final concentration to 20 mM to ensure it is above the critical micelle concentration (CMC) of DTAB which is ~15 mM. The total volume was 83 µL and was adjusted by addition of 1X buffer H to achieve a final DNA and lipid concentration of 300 nM and 135 µM respectively. Samples were incubated at 30 °C (above the phase transition temperature of DMPC which is 25 °C) overnight followed by detergent removal using Pierce™ detergent removal spin columns (cat. no. 87777, Thermo Scientific) according to the manufacturer's protocol. Briefly, the spin columns were spun down at 1500 x *g* for 1 min to remove storage buffer. Columns were equilibrated by three additions of 400 µL of 1X buffer H followed by centrifugation at 1500 x *g* for 1 min. Next, the sample was added, incubated for 2 min and centrifuged at 1500 x *g* for 2 min to retrieve the detergent-free PEG-DLNs in 83 µL of 1X buffer H.

**Density-gradient ultracentrifugation.** Optiprep density gradient medium (60 % iodixanol) was used to prepare different gradients ranging from 2 % to 26 % in 1X buffer H.

| Percentage | Optiprep medium (μL) | 10X buffer H (μL) | Water (μL) |
|---|---|---|---|
| 2 | 16 | 48 | 416 |
| 6 | 48 | 48 | 384 |
| 10 | 80 | 48 | 352 |
| 14 | 112 | 48 | 320 |
| 18 | 144 | 48 | 288 |
| 22 | 176 | 48 | 256 |
| 26 | 208 | 48 | 224 |

83 μL of the detergent-free PEG-DLNs were mixed with 83 μL of 55 % iodixanol to prepare a 27 % iodixanol solution. 0.8 μL open-top thin wall ultracentrifuge tubes (Beckman Coulter Life Sciences, cat. no. 344090) were used for this experiment. 166 μL of the sample + iodixanol (27 % solution) was added to the tube followed by layering with 70 μL of 26 %, 22 %, 18 %, 14 %, 10 %, 6 % and 2 % solutions respectively. The tube was placed with its adapters (Beckman Coulter Life Sciences, cat. no. 356860) in a SW55Ti swinging bucket rotor (Beckman Coulter Life Sciences, cat. no. 342194). Samples were spun at 45000 rpm (~245,000 *g*) for 6 h at RT after which the contents were fractionated into 96-well plates, followed by analysis of lipid signals using fluorescence and DNA by native-SDS PAGE respectively. Fractionated samples were imaged on a GE Typhoon FLE 9500 gel scanner using a 473 nm excitation laser and 510 nm long pass emission filter (ex/em maxima for TfPC lipid). The photomultiplier tube gain was set to 500 and the pixel size was 50 μm. Fluorescence signals were analyzed by the Fiji ImageJ image analysis software.

**Transmembrane peptide incorporation.** The transmembrane (TM) domain of synaptobrevin was modified to contain a biotin at the C-terminal end. The 23-amino acid peptide contains no charged amino acids (IILGVISAIILIIIIVYFSTGSS-Biotin, average molecular weight = 2674.4 gmol$^{-1}$). The C-Terminal "GSS" sequence is an alpha helix breaker and was added to provide flexibility. The peptide was solubilized to a final concentration of 50 μM in an aqueous 100 μM DTAB solution. 8.53 μL (25 pmol) of DNA-PEG-minicircles in 1X buffer H were mixed with 5.62 μL (11.25 μmol) of lipids and 1 μL (50 pmol) of peptide solution. This results in a 1: 450 ratio of DNA: lipid and 1: 2 ratio of DNA: peptide. To this solution, 13.23 μL of 100 mM DTAB were added to adjust the final concentration to 20 mM to ensure it is above the critical micelle concentration (CMC) of DTAB which is ~15 mM. Then, 2 μL of 1 μM QD-streptavidin conjugate were added and mixed well. An excess of the QD-streptavidin could not be added due to low concentration of the commercially available conjugate. The total volume was 83 µL and was adjusted by addition of 1X buffer H to achieve a final concentration of 300 nM DNA and 135 µM of lipids. Samples were incubated at 30 °C (above the phase transition temperature of DMPC which is 25 °C) overnight followed by detergent removal using Pierce™ detergent removal spin columns (cat. no. 87777, Thermo Scientific) according to the manufacturer's protocol to retrieve detergent-free PEG-DLNs with a TM peptide bound to QD-streptavidin.

After ultracentrifugation, samples were fractionated and imaged on a GE Typhoon FLE 9500 gel scanner using a 473 nm excitation laser with a 510 nm long pass emission filter (ex/em maxima for TfPC lipid) and a 635 nm excitation laser with a red long pass emission filter (ex/em maxima for the QD conjugate). The photomultiplier tube gain was set to 500 and the pixel size was 50 µm. Fluorescence signals were analyzed by the Fiji ImageJ image analysis software.

**Synthesis of biotinylated transmembrane peptide**. The peptide $NH_2$-IILGVISAIILIIIIVYFSTGSS-Biotin (MW= 2632.45 g/mol) was synthesized using microwave-assisted solid-phase peptide synthesis (SPPS) on a CEM Liberty Blue 2.0 instrument employing Fmoc-chemistry. Biotin Nova Tag™ resin was used as the solid support, with amino acid activation carried out using N,N'-Diisopropylcarbodiimide and Oxyma in DMF, and piperidine as the base. Resin cleavage and sidechain deprotection were performed using a mixture of trifluoroacetic acid (87.5% v/v), phenol (5% v/v), triisopropyl silane (2.5% v/v), and water (5% v/v) for 2.5 h at room temperature (21 °C). The crude peptide was then precipitated in diethyl ether and collected by vacuum filtration.

#### 2. Atomistic Molecular Dynamics Simulations

##### 2.1. General Simulation Methods

All-atom simulations were performed using NAMD 3.0.1[2] and employed the CHARMM36[3,4] force field with CUFIX[5–7] corrections, TIP3P[8] water model, permanent magnesium hexahydrates,[6] and periodic boundary conditions. Hydrogen mass repartitioning[9] and the SHAKE[10] and SETTLE[11] hydrogen constraint algorithms were used, enabling a 4 fs integration time step. Van der Waals and short-range electrostatic forces were evaluated using an 8–12 Å smooth cutoff scheme. Long-range electrostatic interactions were computed using the particle-mesh Ewald method[12] with a 1-Å grid spacing. The temperature was held at 310 K by a Langevin thermostat applied to non-hydrogen atoms with 1 $ps^{-1}$ damping coefficient. The pressure was maintained at 1 atm by a Nosé–Hoover Langevin piston barostat[13,14] with a 2000 fs period and 1000 fs decay. When present, base-pairing hydrogen bonds were reinforced using additional harmonic bonds with 2.85 Å rest length and 100 kcal $mol^{-1}$ $Å^{-2}$ spring constant. The configuration of a system was written to a trajectory file every 10 ps of simulation.

#### 2.2. System Assembly and Simulation of DLN and Bare Lipid Systems

An idealized model of a DNA ring composed of 147-bp of DNA with sequence $(GTGAAAAAGTGTGAAAAAGT)_7$ and containing seven evenly distributed nicks was prepared using a custom script that employed a brief restrained mrDNA[15] simulation to ensure the adenine nucleotides would face the outside of the ring. VMD[16] and the psfgen program were used to combine a 79-Å radius patch of 90% DMPC, 9% DMTAP and 1% DMPE (564 total lipids) cut to fit, without steric clashes, inside the DNA ring functionalized by 3-amine linkers per to PEG per 21-nt ssDNA fragment (described in the experimental methods section), and 80 DTAB molecules arranged near the DNA. The lipid bilayer described above was excised from an idealized patch generated by CHARMM-GUI.[17,18] The entire system was submerged in a 21×21×8.5 $nm^3$ box of 100 mM $Na_2SO_4$ and 6 mM $MgCl_2$ electrolyte, while the system was made neutral by adding ~45% of the solute charge in additional $Mg^{2+}$ ions and removing $Cl^-$ and $SO_4^{2-}$ corresponding to ~5% and of the 50% of the excess charge, respectively.

After assembly, conjugate gradient minimization was applied to the system for 1000 steps, followed by equilibration with harmonic restraints applied to non-hydrogen lipid atoms, excluding DTAB using a 1 kcal $mol^{-1}$ $Å^{-2}$ spring constant that was halved every 16 nanoseconds for seven cycles (112 ns total) until being entirely removed. During the first 120 ns of equilibration (including lipid-restrained equilibration), an mrDNA-generated elastic network of restraints that preserve base pairing and stacking was applied to the DNA, after which only base pair hydrogen bonds were restrained. After 378 ns of simulation, the DLN had developed a significant tilt with respect to the short axis of the system, and a colvars[19] tilt angle variable was created that included every fourth lipid C2 atom, using the initial configuration as a reference. The tilt angle is calculated by decomposing the rotation that best aligns the C2 atoms into two rotations, a spin rotation about the *z*-axis of the system (initially normal to the bilayer), and a tilt rotation in the *xy*-plane. The collective variable was calculated every 100 steps and was used in a harmonic restraint that guided the cosine of the tilt angle from 0.9952 (~5.6°) to 1 over a 10 ns period using a stiff 10,000 kcal $mol^{-1}$ (~3 kcal $mol^{-1}$ $degree^{-2}$) spring constant.

A second DLN system with 0.5 PEG per DNA turn system was constructed as described above, except the functionalized DNA strands contained only the central PEG molecule. The simulations were performed as described above, except the final 12 nanoseconds of restrained equilibration were skipped, and the colvars tilt angle restraint was applied from the start of the simulation. Finally, a complementary "bare lipid" system designed to provide a comparison without the DNA scaffold was constructed using identical protocols as described above, except that the functionalized DNA was not added to the system, and the charge of the system was neutralized by adding excess $SO_4^{2-}$. The simulation protocols were identical to the DLN system, except that the absence of DNA in the system ensured that no contacts could be made between periodic images of the bilayer, even without colvars tilt angle restraints.

##### 2.3. Analysis of MD Trajectories

Analysis of the simulations was performed with VMD and MDAnalysis using preprocessed trajectories where the coordinates of the system were transformed to place the center of mass of the lipid bilayer C2 atoms (representative of the headgroup) at the origin and align their principal axes with cartesian axes, simplifying subsequent calculations.

The occluded fraction of solvent accessible surface area of the lipid tails was estimated using the `measure sasa` feature of VMD with a 2.0 Å probe radius to compute the accessible area of the lipid tails with and without considering PEG as part of the solvent. Subtracting the areas provides the lipid tail area occluded by the PEG and DTAB, and dividing by the area that treats PEG as part of the solvent provides the occluded fraction solvent accessible lipid tail area. The analysis of contacts was performed using the `measure contacts` feature of VMD with a 4 Å cutoff to consider two atoms in contact.

The fraction of DMTAP or DTAB in a given region was computed by counting the number of C2 atoms of the molecule of interest found within the region, and dividing by the total number of lipid (plus DTAB) C2 atoms found in the region. The regions were defined by bisecting the bilayer into two leaflets and according to proximity to the bilayer edge for a given configuration according to the following procedure. The `measure volinterior` feature of VMD with an isovalue of 0.01 and resolution of 6 to extract a molecular surface of all lipid atoms (excluding DTAB), marking voxels as inside or outside the surface. Subsequently, a distance from the edge was calculated for each molecule using distance projected into the plane of the bilayer of its C2 atom from the center of the nearest pixel that was, on average over the normal axis of a 2-nm thick slab centered on the bilayer and sharing its normal, outside the bilayer. Except where specified, characterization of bilayer properties described below involved analysis of only those lipids with C2 atom contained within a 4 nm radius cylinder centered on the bilayer with axis along the bilayer normal. For a given configuration, the thickness of the bilayer was assessed by computing the density profile along the bilayer normal in 0.5 Å bins and fitting a gaussian distribution to the density for each leaflet. The area pear lipid headgroup was determined by counting the C2 atoms in the 4-nm radius cylinder. The curvature was estimated by performing a nonlinear least squares fit of a sphere to the lipid headgroup coordinates of each leaflet with shift along the bilayer normal and curvature taken as independent variables; the result obtained for the two leaflets were averaged. The number of lipid headgroups at the edge of each system was computed as the number of lipid C2 atoms in the 1-nm thick slab centered along the bilayer and sharing the bilayer normal vector with no restriction on which lipids are considered for the analysis.

#### Supporting Figures

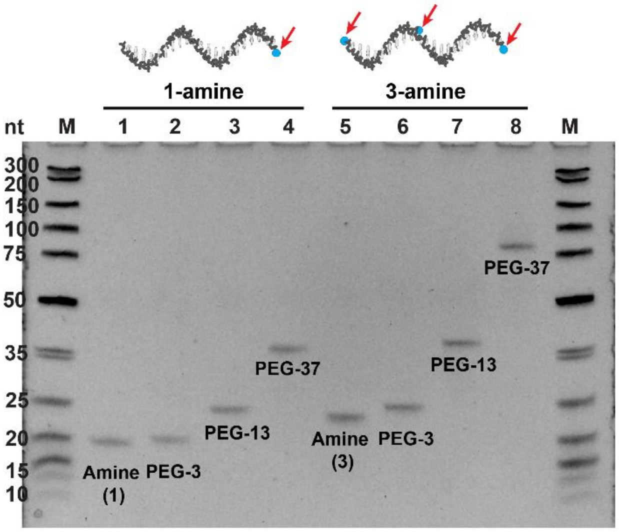


**Figure S1:** Denaturing gel electropherogram of PEGylated oligonucleotides. M: Ultra-low range DNA ladder. Lanes 1-4: 21-base oligonucleotide with one amine modification before and after reaction with PEG-3, PEG-13 and PEG-37 NHS esters respectively. Lanes 5-8: 21-base oligonucleotide containing three amine modifications before and after reaction with PEG-3, PEG-13 and PEG-37 NHS esters respectively.

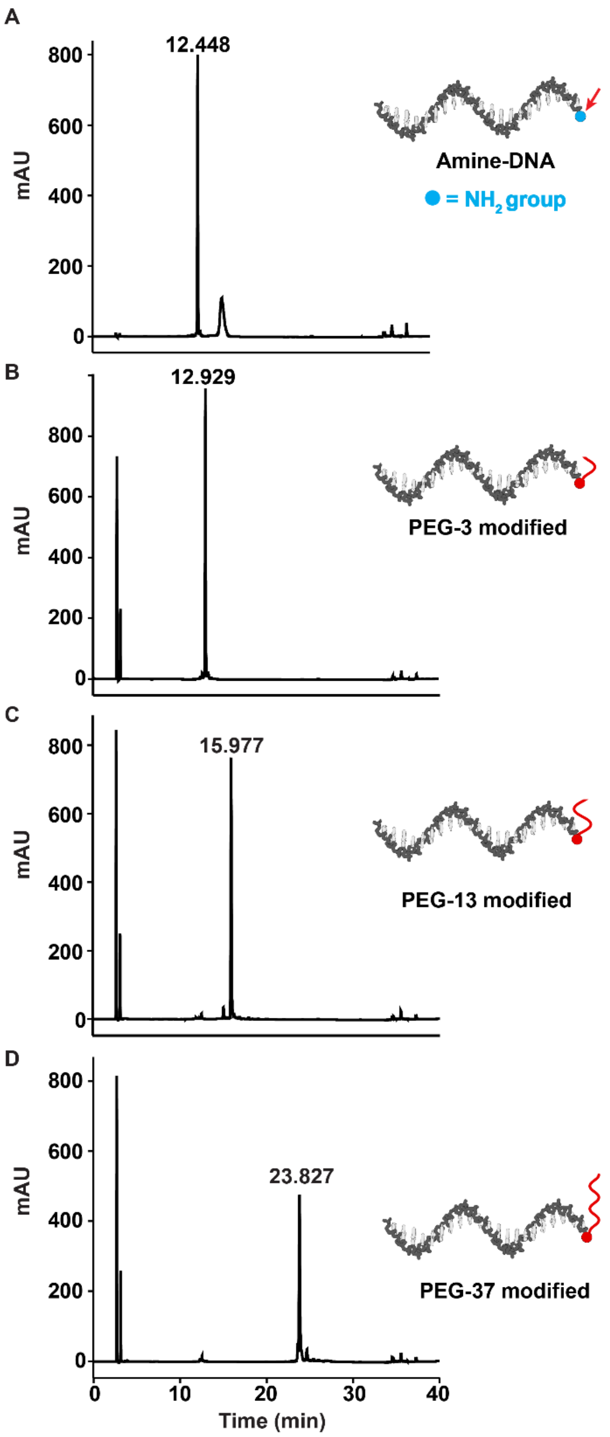


**Figure S2.** HPLC chromatograms of PEGylated oligonucleotides. (A) HPLC chromatogram of starting 1 amine oligonucleotide with retention time ~12 min. 1 amine oligonucleotide coupled to (B) PEG-3 NHS ester with a shift in retention time to ~13 min, (C) PEG-13 NHS ester with a shift in retention time to ~16 min, (D) PEG-37 NHS ester with a shift in retention time to ~24 min. Increasing shifts in retention time are attributed to the increasing hydrophobicity of the PEG chain lengths. Gradient: 3% to 30% acetonitrile in 30 min; 30% to 100% acetonitrile in the next 5 min; column wash with 100% acetonitrile for 3 min.

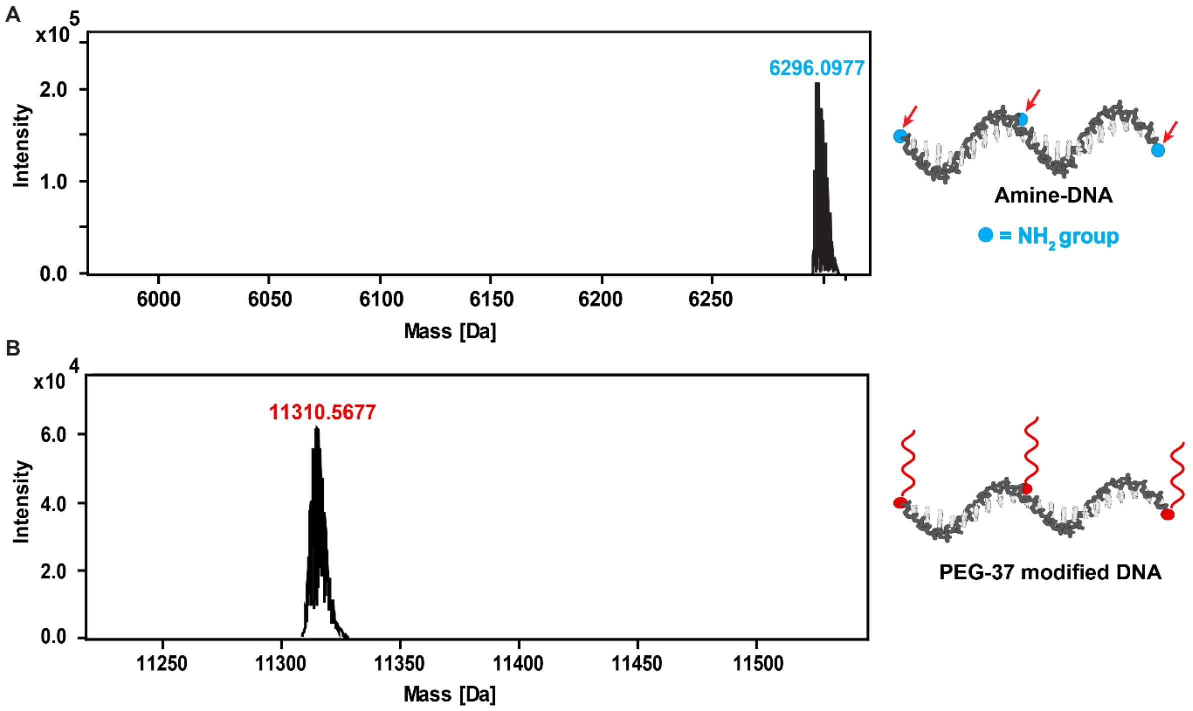


**Figure S3**. Mass spectra. Deconvoluted high-resolution mass spectra after HPLC-ESI-MS: (A) 3-amine oligonucleotide, expected average mass = 6298.4 $gmol^{-1}$, (B) 3 PEG-37-oligonucleotide conjugate, expected average mass = 11311.43 $gmol^{-1}$. Note that the peak pattern comes from the natural isotope distribution and only the highest peak is labeled for clarity.

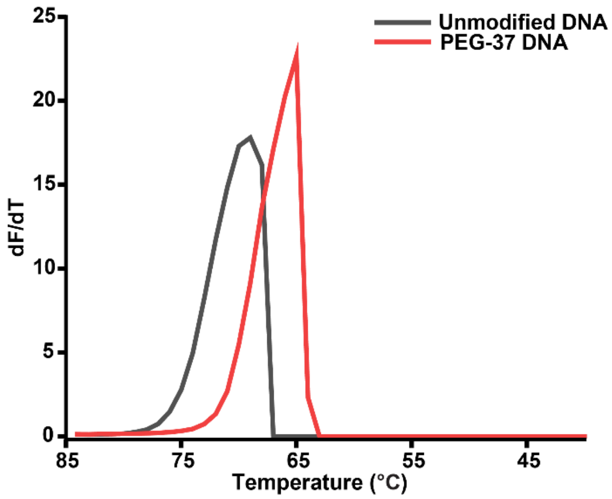


**Figure S4.** $T_m$ analysis for PEGylated DNA. Melting temperature ($T_m$) obtained by plotting first derivative of change in fluorescence with respect to temperature against change in temperature (dF/dT). Black: DNA without amine group ($T_m$ = 69 °C). Red: DNA with 3 PEG-37 modifications ($T_m$ = 65 °C).

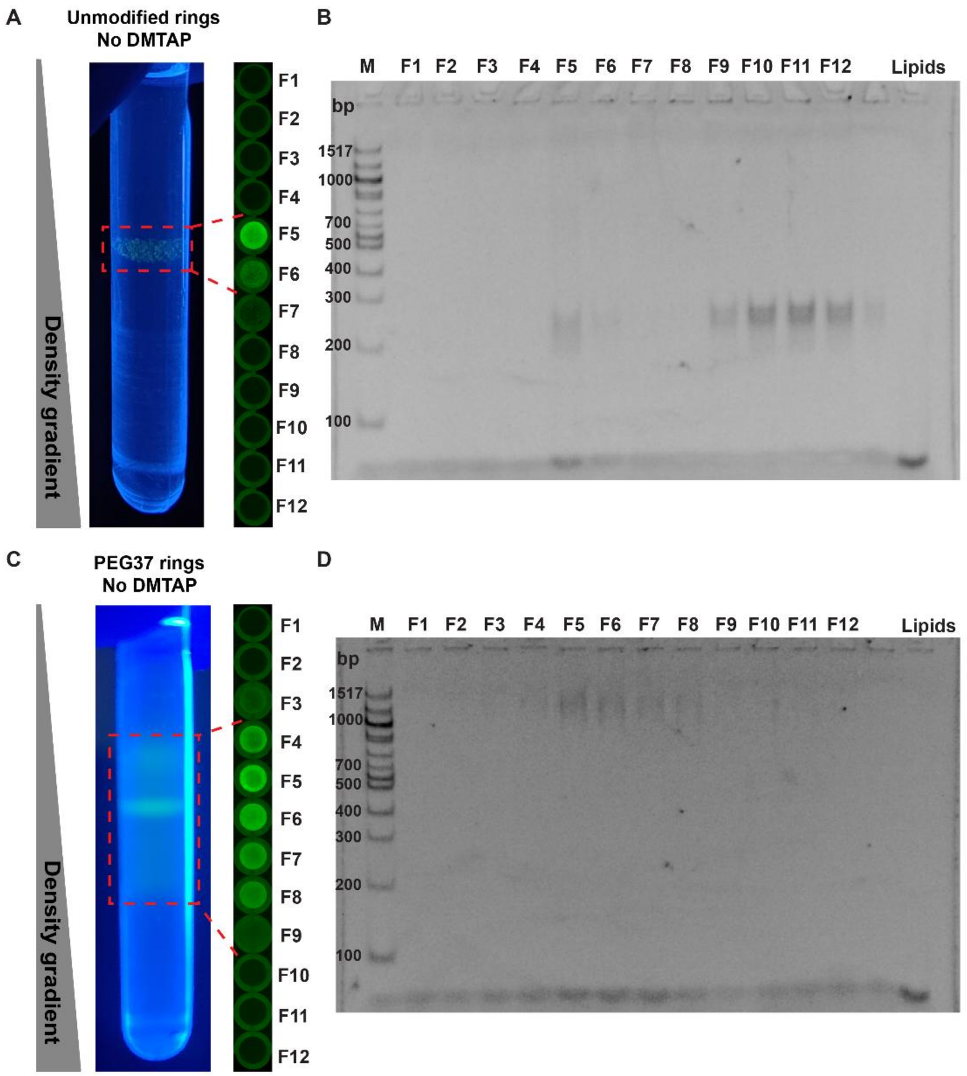


**Figure S5**. Reconstitution in the absence of cationic lipid. DLN synthesis attempts with ds-minicircles and DMPC, TfPC without any DMTAP. A-B: reconstitution with unmodified ds-minicircles, C-D: reconstitution with PEG-ds-minicircles. (A-B) Fractions 9-12 show most ds-minicircles do not colocalize with lipids except for F5-F6 which show colocalization due to the formation of an electrostatic aggregate . In C-D, ds-minicircles are observed in F4-F10 and colocalize with lipids. However, the fluorescence is distributed over several fractions and does not appear concentrated as in samples with DMTAP. See Figure S8 for additional experimental details.

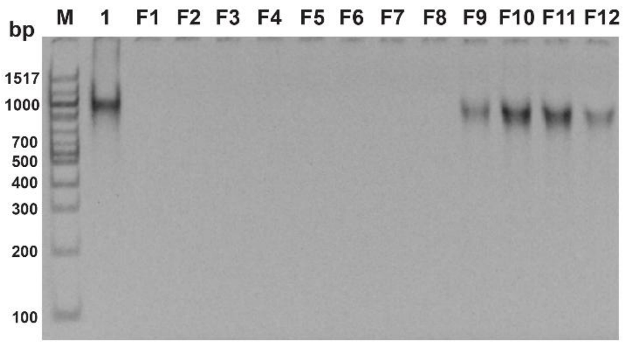


**Figure S6.** Gel electropherogram of a PAGE gel containing SDS and $MgCl_2$ of the fractions collected after isopycnic ultracentrifugation showing the ds-PEG-minicircles in the bottom fractions (F9-F12). M: 100 bp dsDNA ladder. Lane 1: ds-PEG-minicircle (control).

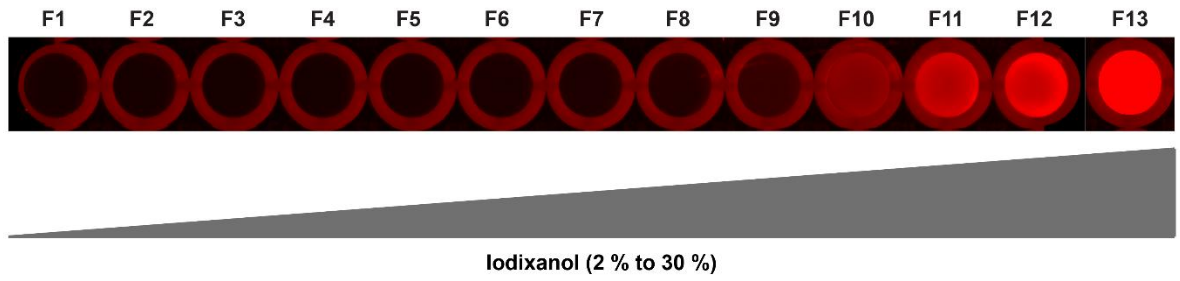


**Figure S7.** Non-specific interactions of Nile red (lipid stain) with iodixanol (density gradient medium). Fractions F1-F13 contain increasing amounts of iodixanol (in buffer) ranging from 2-30 %, fractionated after isopycnic ultracentrifugation. A constant amount of the Nile red was added to each well and imaged in a fluorescence scanner with a 532 nm excitation laser. This demonstrates that lipid bilayers cannot selectively be detected with Nile red due to the non-specific staining with iodixanol.

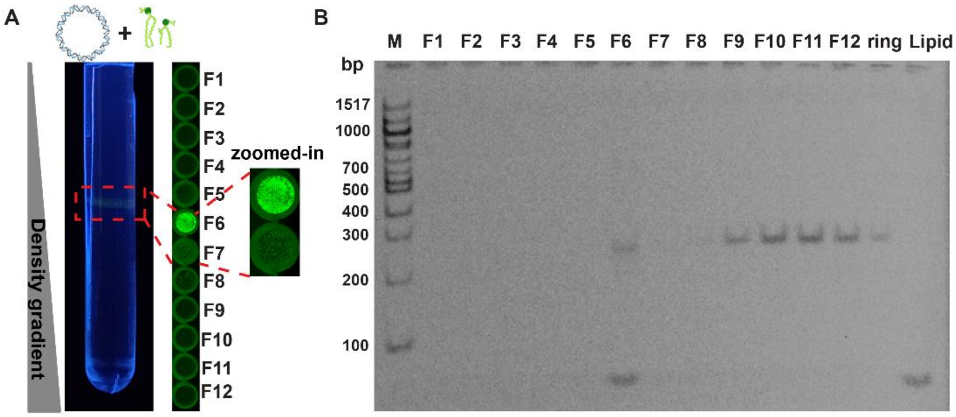


**Figure S8.** Control experiment without PEG modifications. Analysis of a DLNs experiment with unmodified ds-minicircles (no PEG modification). The lipid formulation contained 1% fluorescent lipid (top fluor-PC). (A) The photograph of the tube after ultracentrifugation was taken under UV illumination to visualize fluorescent lipids. The content of the ultracentrifugation tube (left) was fractionated into the wells of a 96-well plate and imaged in a fluorescence scanner (right), indicating lipids in the 6$^{th}$ and 7$^{th}$ fraction (F6-F7). Note that the lipids appear visually aggregated in the zoomed-in fluorescence scan, indicating that nanodiscs are unstable. (B) PAGE analysis of the fractions taken from (A). The gel contained SDS to solubilize lipids and $MgCl_2$ to stabilize dsDNA and stained with Sybr Gold to detect DNA (band comigrating with the 300 bp band of the ladder). The majority of the ds minicircles was found in fractions 9-12, not colocalized with the lipid signal (lower band in F6). The small fraction of the minicircles in F6 is possibly due to the formation of an electrostatic complex. M: 100 bp dsDNA ladder. Ring: Unmodified ds-minicircle (control); control lipids were solubilized in SDS and form mixed micelles migrating a little faster than the 100 bp marker lane.

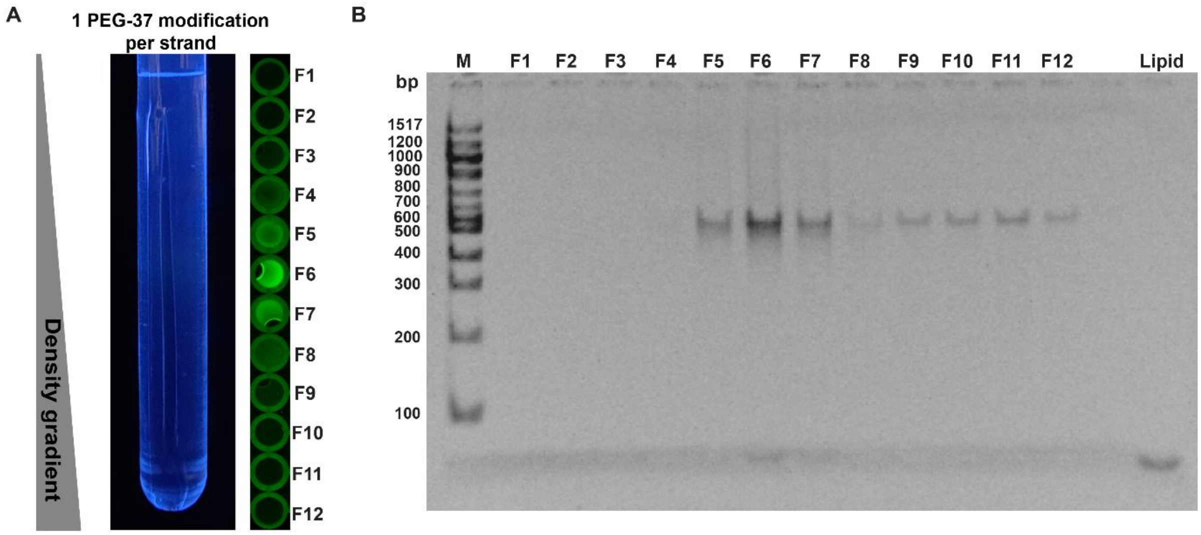


**Figure S9.** Reducing the number of PEG modifications. DLNs synthesis is inefficient with ds-PEG-minicircles containing only one PEG-37 modification per strand, or seven PEG-37 modifications per ds-ring instead of 21. (A) No sharp band is observed (A) and instead lipids are spread around fractions F5-F7, while ds minicircles appear in fractions F5-F12 (B). See Figure S8 for additional experimental details.

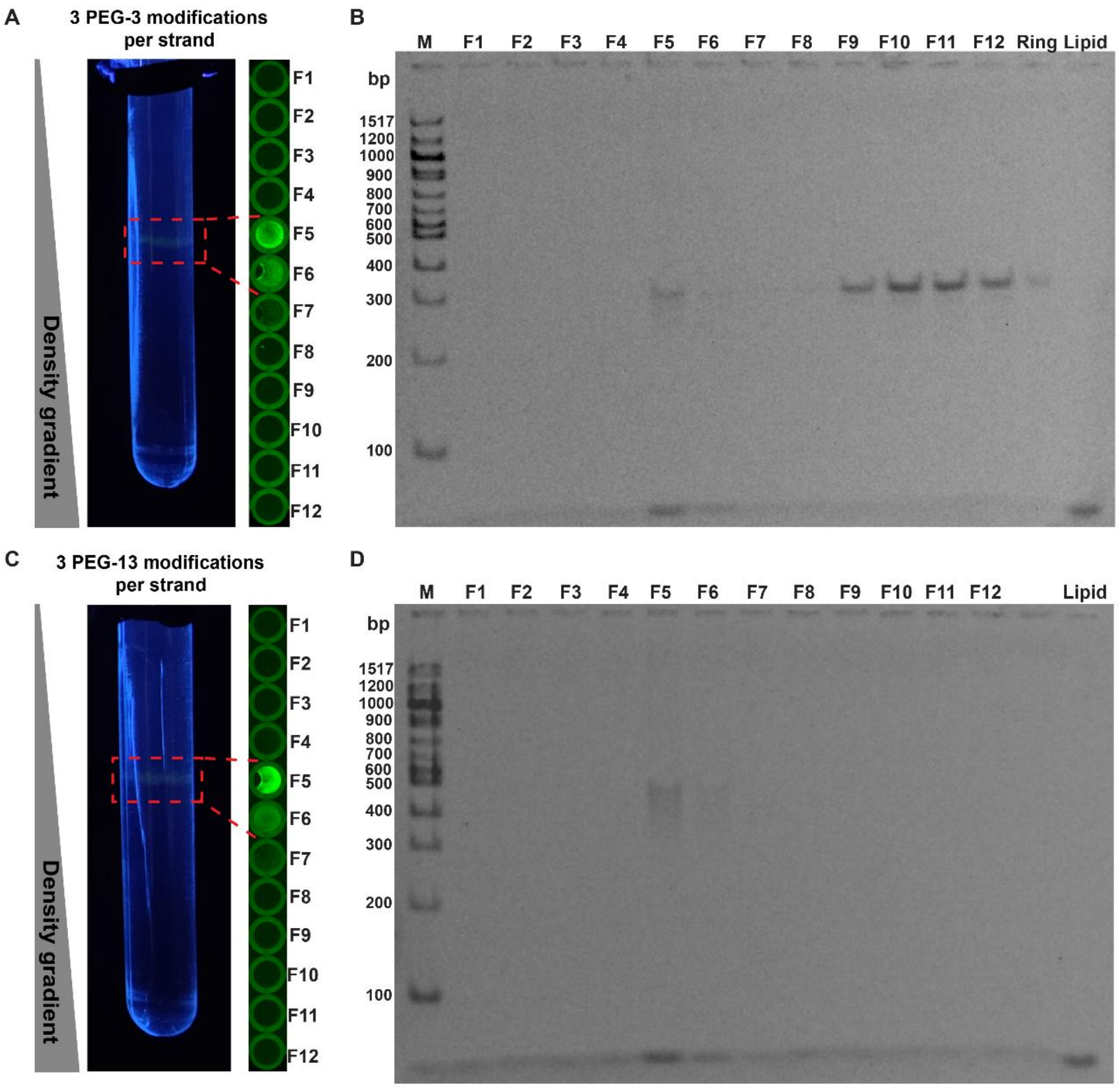


**Figure S10.** Shortening PEG chains. DLNs are not efficiently formed with ds-minicircles with shorter PEG modifications. ds-Minicircles contained 21 PEG-3 (A-B; three ethylene glycol repeats) or 21 PEG-13 modifications per ds-ring (C-D) instead of the optimized PEG-37 chain length. In both cases, lipids formed a fuzzy bands around F5-6, but only a fraction of DNA (upper band) colocalized with the lipids. The overall intensity is low indicating significant loss of sample in the detergent removal step. See Figure S8 for additional experimental details.

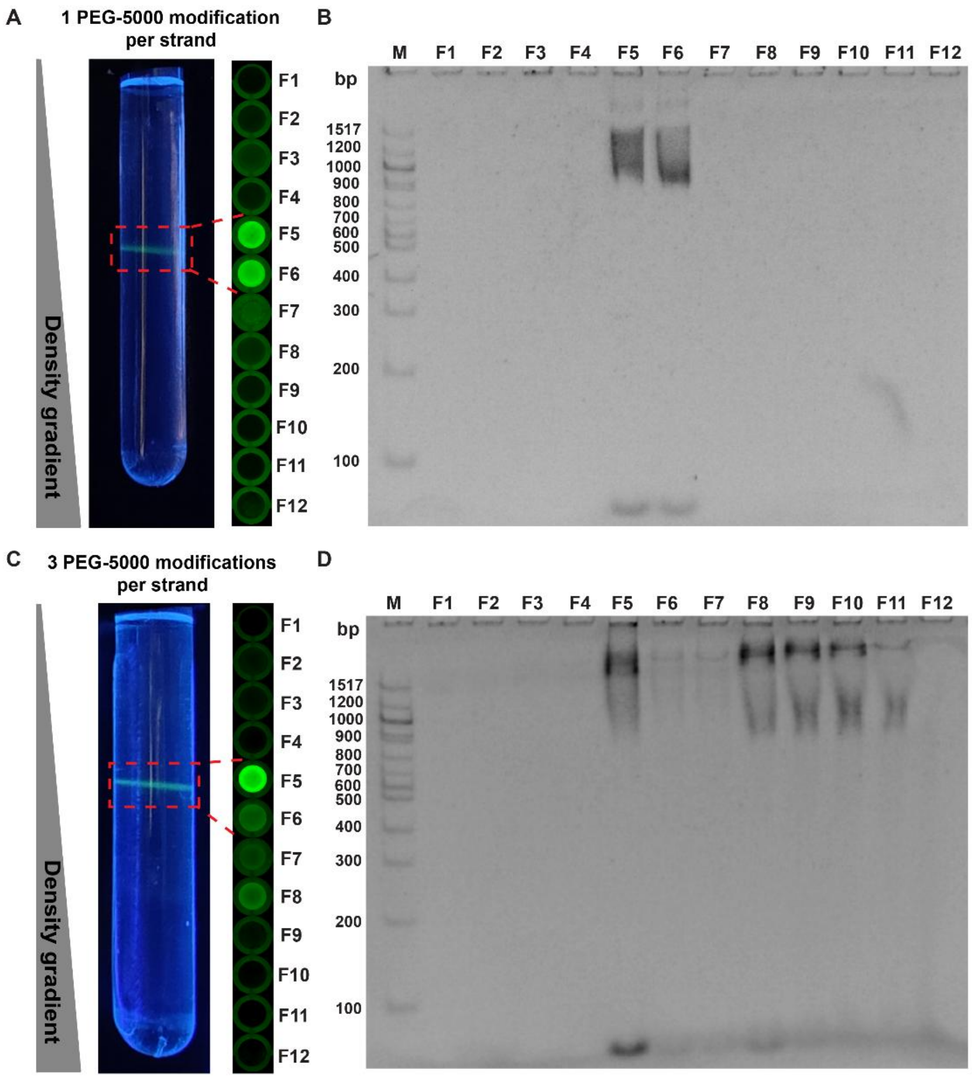


**Figure S11.** Increasing the PEG chain length. DLN synthesis attempts with ds-minicircles with longer PEG-5000 (~5 kDa) instead of PEG-37 (~1.7 kDa) modifications. A-B: one modification per strand or seven PEG chains per ds-minicircle, C-D: three modifications per strand or 21 per ds-minicircle. (A-B) Fractions 5-6 show most ds-minicircles colocalized with lipids indicating successful DLN formation. However, in C-D, some ds minicircles (F8-10) do not colocalize with lipids suggesting that extensive modification with long PEG chains hinders DLN formation. Note that unlike the atomically precise PEG-37, PEG-5000 is a polydisperse polymer and therefore DNA bands are more smeared out. See Figure S8 for additional experimental details.

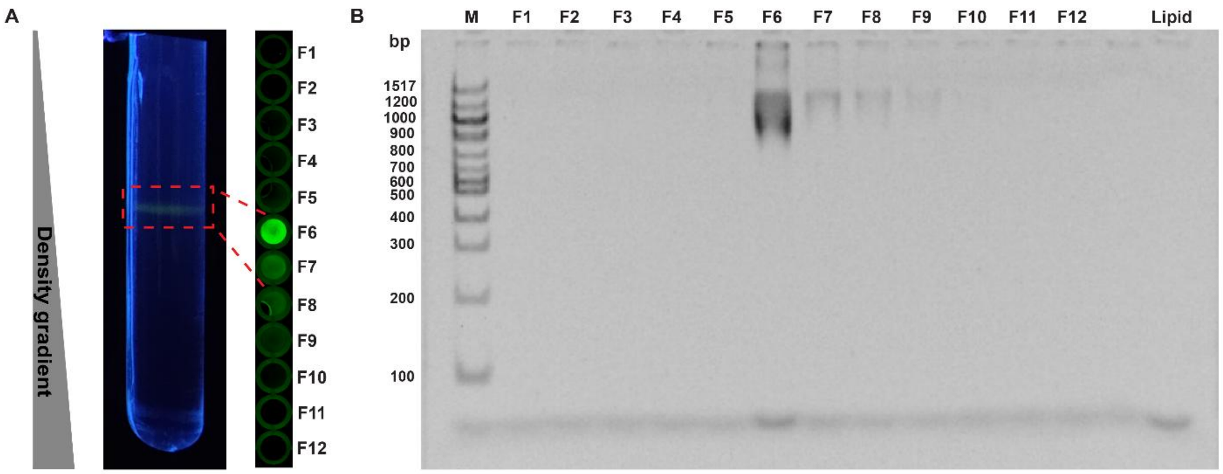


**Figure S12.** Increasing minicircle flexibility. In this experiment, were designed ds-PEG-minicircles with 21 PEG-37 modifications to have a two-nucleotide single-stranded gap between all seven oligonucleotides on ss scaffold ring. While lipids colocalized with the ds minicircles indicating successful DLN formation, the results did not show any improvement over the design without gaps and was therefore discontinued. See Figure S8 for additional experimental details.

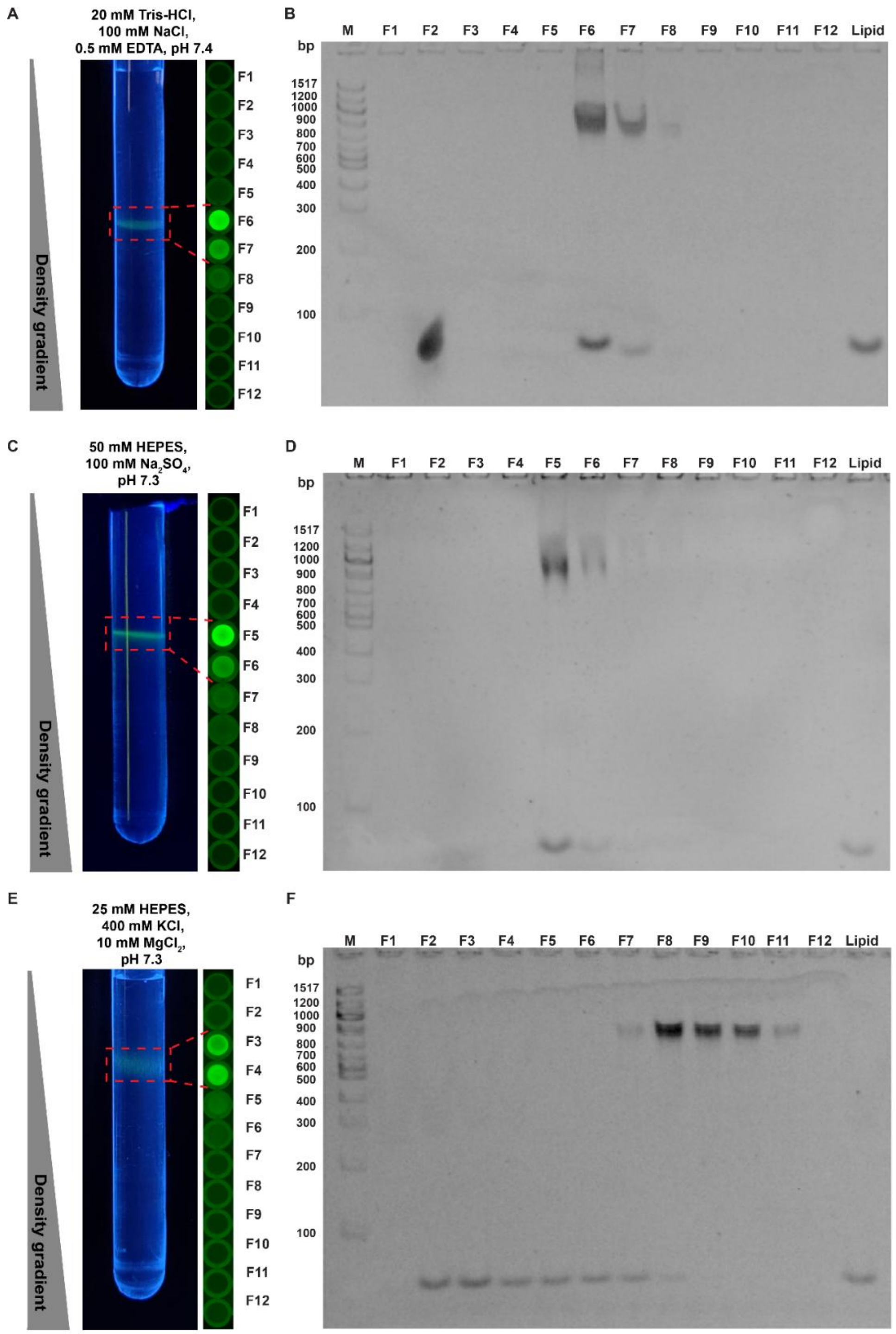


**Figure S13.** Testing buffer conditions. Analysis of DLNs formed with ds-PEG-minicircles (21 x PEG-37) under different buffer conditions. A-D: DLNs form in common buffers containing 100 mM sodium salts as indicated by the colocalization of lipids (bottom band) and DNA (upper band) in fractions 5-7. E-F: Lipids and DNA are not colocalized in the same fractions indicating that DLNs do not form in a high-salt buffers containing 400 mM $K^+$ and 10 mM $Mg^{2+}$ or that they are unstable . See Figure S8 for additional experimental details.

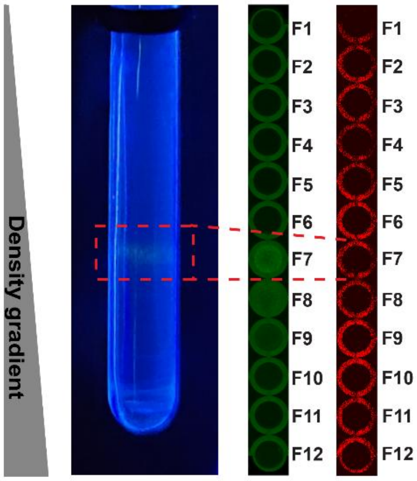


**Figure S14.** Transmembrane peptide reconstitution without PEG. Analysis of a nanodisc reconstitution experiment with lipids, biotinylated TMD peptide and streptavidin-modified quantum dots (QD), but without PEGs on the ds-minicircles. A grainy green fluorescing band in fraction 7 contained lipids in an aggregated state, but no red fluorescing QDs (right wells). In this experiment, QDs were presumably removed along with the peptides in the detergent removal column. This demonstrates that only nanodiscs with a shielding DNA rim can pass the detergent removal column. See Figure S8 for additional experimental details.

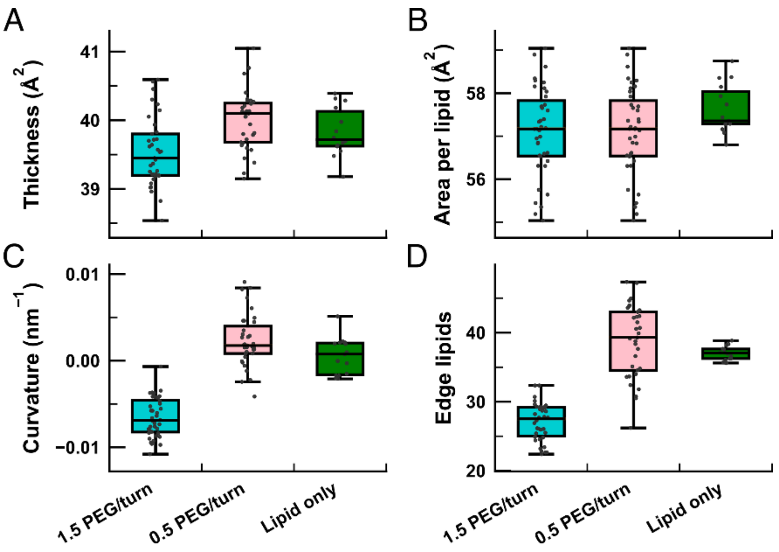


**Figure S15.** Characterization of lipid bilayers in DLNs with 1.5 (cyan) and 0.5 PEG per DNA turn (pink), as well as a lipid-only bilayer (green). Lipid bilayer properties of the central 4-nm radius region of each bilayer, including bilayer thickness (A), area per lipid (B) and curvature (C). (D) Number of lipid headgroups observed along the edges of a DLN and bare bilayer. Here, a lipid is defined to lay along the edge of the bilayer if its C2 atom is located within a 1 nm slab centered on and aligned with the bilayer.

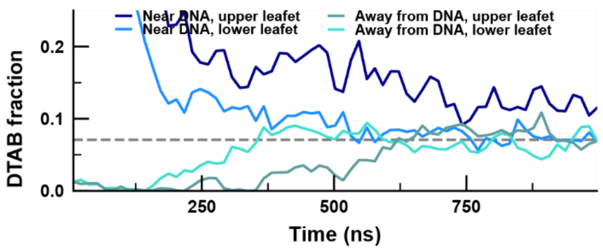


**Figure S16.** DTAB analysis. An analysis of DTAB localization in the lipid bilayer following the same method used for DMTAP (Figure 3). In the beginning of the simulation, all DTAB molecules were placed at the interface between DNA and lipids. The simulation produces only a modest DTAB enrichment in the upper leaflet near the DNA.

#### Captions for movies

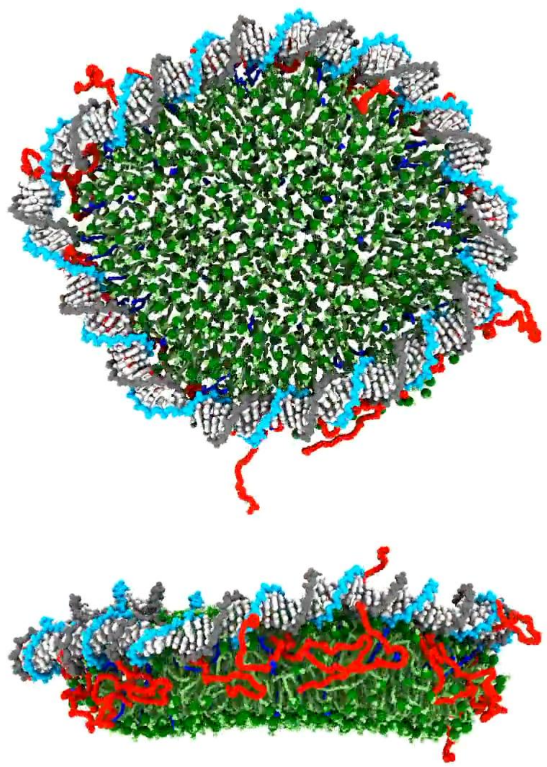

**Movie 1.** All-atom molecular dynamics simulations of PEGylated DLNs. The movie includes the restrained equilibration lasting ~120 ns and subsequent unrestrained simulation lasting nearly 900 ns. A DNA construct (cyan and gray) functionalized with PEG molecules (red) stabilize the lipid bilayer (DMPC, DMTAP and DMPE; green with C2 atom depicted as dark green sphere) doped with DTAP detergent molecules (blue). The top image depicts the DLN from above and the bottom image depicts the same system rotated by 90°.

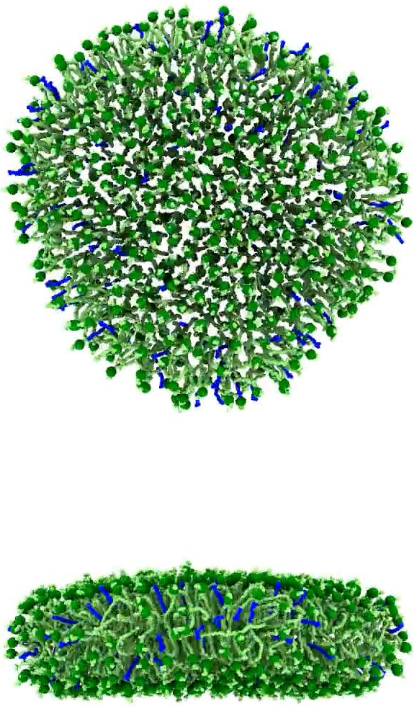

**Movie 2.** All-atom molecular dynamics simulations of a bare lipid bilayer. The movie includes the restrained equilibration lasting ~120 ns and subsequent unrestrained simulation lasting ~400 ns. The system is depicted using the same representations as in **Movie 1**. Note that such a system would be experimentally unstable due to fusion of multiple of these patches into liposomes.

### References

(1) Iric, K.; Subramanian, M.; Oertel, J.; Agarwal, N. P.; Matthies, M.; Periole, X.; Sakmar, T. P.; Huber, T.; Fahmy, K.; Schmidt, T. L. DNA-Encircled Lipid Bilayers. *Nanoscale* **2018**, *10* (39), 18463–18467. https://doi.org/10.1039/C8NR06505E.

(2) Phillips, J. C.; Hardy, D. J.; Maia, J. D. C.; Stone, J. E.; Ribeiro, J. V.; Bernardi, R. C.; Buch, R.; Fiorin, G.; Hénin, J.; Jiang, W.; McGreevy, R.; Melo, M. C. R.; Radak, B. K.; Skeel, R. D.; Singharoy, A.; Wang, Y.; Roux, B.; Aksimentiev, A.; Luthey-Schulten, Z.; Kalé, L. V.; Schulten, K.; Chipot, C.; Tajkhorshid, E. Scalable Molecular Dynamics on CPU and GPU Architectures with NAMD. *J. Chem. Phys.* **2020**, *153* (4), 044130. https://doi.org/10.1063/5.0014475.

(3) Hart, K.; Foloppe, N.; Baker, C. M.; Denning, E. J.; Nilsson, L.; MacKerell, A. D. Jr. Optimization of the CHARMM Additive Force Field for DNA: Improved Treatment of the BI/BII Conformational Equilibrium. *J. Chem. Theory Comput.* **2012**, *8* (1), 348–362. https://doi.org/10.1021/ct200723y.

(4) Klauda, J. B.; Venable, R. M.; Freites, J. A.; O'Connor, J. W.; Tobias, D. J.; Mondragon-Ramirez, C.; Vorobyov, I.; MacKerell, A. D. Jr.; Pastor, R. W. Update of the CHARMM All-Atom Additive Force Field for Lipids: Validation on Six Lipid Types. *J. Phys. Chem. B* **2010**, *114* (23), 7830–7843. https://doi.org/10.1021/jp101759q.

(5) Yoo, J.; Aksimentiev, A. New Tricks for Old Dogs: Improving the Accuracy of Biomolecular Force Fields by Pair-Specific Corrections to Non-Bonded Interactions. *Phys. Chem. Chem. Phys.* **2018**, *20* (13), 8432–8449. https://doi.org/10.1039/C7CP08185E.

(6) Yoo, J.; Aksimentiev, A. Improved Parametrization of Li+, Na+, K+, and Mg2+ Ions for All-Atom Molecular Dynamics Simulations of Nucleic Acid Systems. *J. Phys. Chem. Lett.* **2012**, *3* (1), 45–50. https://doi.org/10.1021/jz201501a.

(7) Yoo, J.; Aksimentiev, A. Improved Parameterization of Amine–Carboxylate and Amine–Phosphate Interactions for Molecular Dynamics Simulations Using the CHARMM and AMBER Force Fields. *J. Chem. Theory Comput.* **2016**, *12* (1), 430–443. https://doi.org/10.1021/acs.jctc.5b00967.

(8) Jorgensen, W. L.; Chandrasekhar, J.; Madura, J. D.; Impey, R. W.; Klein, M. L. Comparison of Simple Potential Functions for Simulating Liquid Water. *J. Chem. Phys.* **1983**, *79* (2), 926–935. https://doi.org/10.1063/1.445869.

(9) Balusek, C.; Hwang, H.; Lau, C. H.; Lundquist, K.; Hazel, A.; Pavlova, A.; Lynch, D. L.; Reggio, P. H.; Wang, Y.; Gumbart, J. C. Accelerating Membrane Simulations with Hydrogen Mass Repartitioning. *J. Chem. Theory Comput.* **2019**, *15* (8), 4673–4686. https://doi.org/10.1021/acs.jctc.9b00160.

(10) Ryckaert, J.-P.; Ciccotti, G.; Berendsen, H. J. C. Numerical Integration of the Cartesian Equations of Motion of a System with Constraints: Molecular Dynamics of *n*-Alkanes. *Journal of Computational Physics* **1977**, *23* (3), 327–341. https://doi.org/10.1016/0021-9991(77)90098-5.

(11) Miyamoto, S.; Kollman, P. A. Settle: An Analytical Version of the SHAKE and RATTLE Algorithm for Rigid Water Models. *Journal of Computational Chemistry* **1992**, *13* (8), 952–962. https://doi.org/10.1002/jcc.540130805.

(12) Darden, T.; York, D.; Pedersen, L. Particle Mesh Ewald: An N·log(N) Method for Ewald Sums in Large Systems. *J. Chem. Phys.* **1993**, *98* (12), 10089–10092. https://doi.org/10.1063/1.464397.

(13) Feller, S. E.; Zhang, Y.; Pastor, R. W.; Brooks, B. R. Constant Pressure Molecular Dynamics Simulation: The Langevin Piston Method. *J. Chem. Phys.* **1995**, *103* (11), 4613–4621. https://doi.org/10.1063/1.470648.

(14) Martyna, G. J.; Tobias, D. J.; Klein, M. L. Constant Pressure Molecular Dynamics Algorithms. *J. Chem. Phys.* **1994**, *101* (5), 4177–4189. https://doi.org/10.1063/1.467468.

(15) Maffeo, C.; Aksimentiev, A. MrDNA: A Multi-Resolution Model for Predicting the Structure and Dynamics of DNA Systems. *Nucleic Acids Research* **2020**, *48* (9), 5135–5146. https://doi.org/10.1093/nar/gkaa200.

(16) Humphrey, W.; Dalke, A.; Schulten, K. VMD: Visual Molecular Dynamics. *Journal of Molecular Graphics* **1996**, *14* (1), 33–38. https://doi.org/10.1016/0263-7855(96)00018-5.

(17) Jo, S.; Kim, T.; Iyer, V. G.; Im, W. CHARMM-GUI: A Web-Based Graphical User Interface for CHARMM. *Journal of Computational Chemistry* **2008**, *29* (11), 1859–1865. https://doi.org/10.1002/jcc.20945.

(18) Lee, J.; Cheng, X.; Swails, J. M.; Yeom, M. S.; Eastman, P. K.; Lemkul, J. A.; Wei, S.; Buckner, J.; Jeong, J. C.; Qi, Y.; Jo, S.; Pande, V. S.; Case, D. A.; Brooks, C. L. I.; MacKerell, A. D. Jr.; Klauda, J. B.; Im, W. CHARMM-GUI Input Generator for NAMD, GROMACS, AMBER, OpenMM, and CHARMM/OpenMM Simulations Using the CHARMM36 Additive Force Field. *J. Chem. Theory Comput.* **2016**, *12* (1), 405–413. https://doi.org/10.1021/acs.jctc.5b00935.

(19) Fiorin, G.; Klein, M. L.; Hénin, J. Using Collective Variables to Drive Molecular Dynamics Simulations. *Molecular Physics* **2013**, *111* (22–23), 3345–3362. https://doi.org/10.1080/00268976.2013.813594.
